# Structural variants from donor-specific assemblies affect recurrent chromatin contact divergence in humans

**DOI:** 10.64898/2026.10.05.756789

**Authors:** Lingbin Ni, Jiadong Lin, DongAhn Yoo, Taralynn Mack, Isaac Wong, Julie Wertz, Katherine M. Munson, Kendra Hoekzema, Human Pangenome Reference Consortium (HPRC), Xinghua Shi, Jian Ma, William S. Noble, Evan E. Eichler

## Abstract

GRAPHICAL ABSTRACT

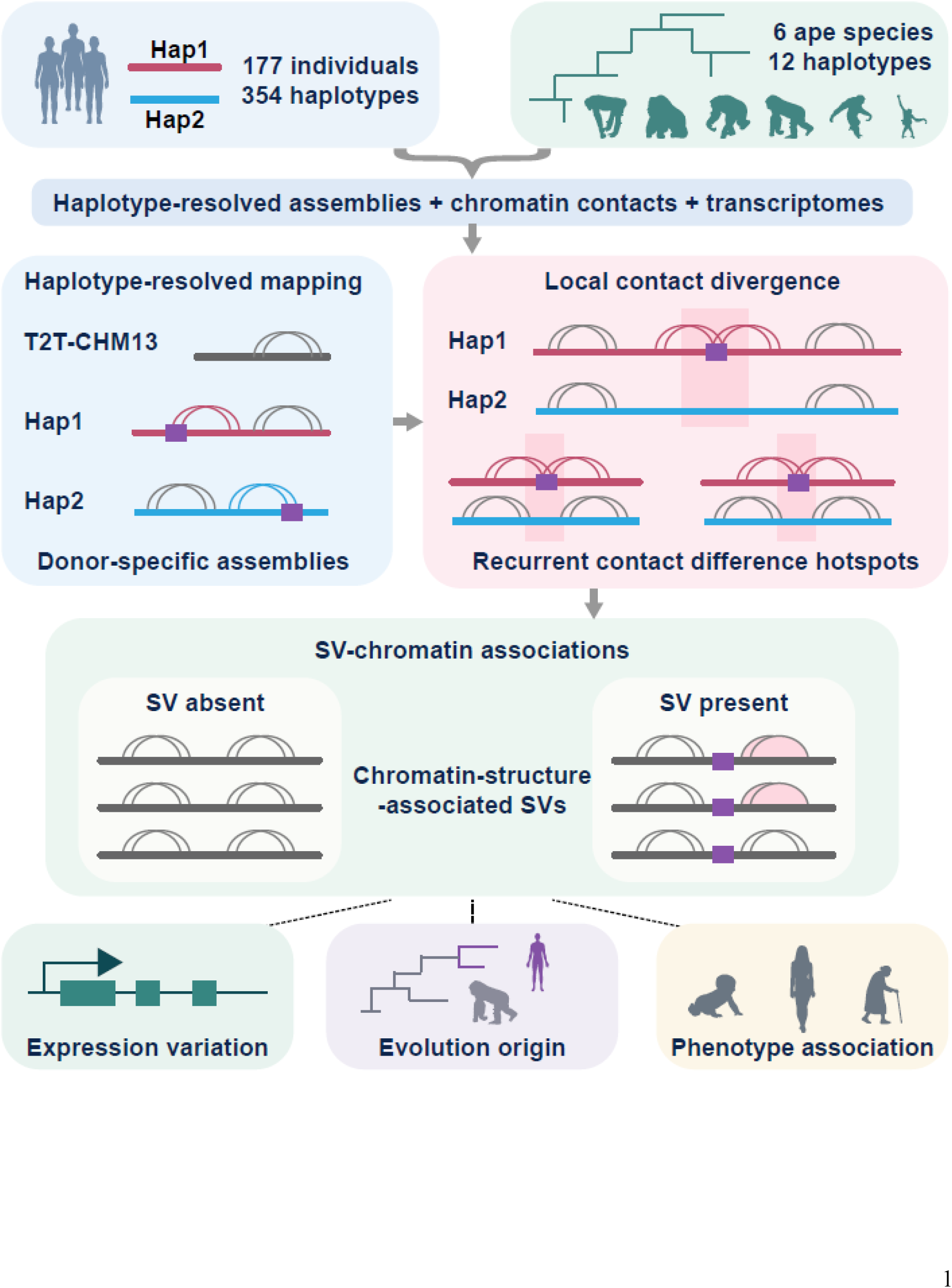

Structural variants (SVs) are a major source of human genetic diversity, yet their contribution to three-dimensional (3D) genome organization remains difficult to resolve using a standard reference genome. Here, we integrate haplotype-resolved chromatin interaction maps across 354 human and 12 nonhuman ape haplotypes with matched genome assemblies and long-read transcriptomes to determine how inherited SVs influence chromatin organization. We observe significant improvements in contact mapping, with 13.2% of cis Hi-C contacts showing >25 kbp changes in inferred genomic separation and an additional 2.8% of valid contact pairs recovered when using donor-specific assemblies instead of standard references. We show that haplotype-specific chromatin differences are highly selective rather than pervasive, concentrating within 1,790 recurrent contact difference hotspots associated with a restricted subset of SVs. We identified 2,495 chromatin-structure–associated SVs that are enriched in regulatory genomic contexts and are linked to expression variation. Notably, more than 90% of these SVs are derived in the human lineage, consistent with recent remodeling of regulatory architecture at the population level. These findings highlight the importance of incorporating haplotype diversity from pangenomes into studies of chromatin organization and provide a framework for understanding how inherited genomic differences are linked to functional consequences across populations and evolution.

## INTRODUCTION

The three-dimensional (3D) organization of the genome is a fundamental layer of genome function that influences transcription, cellular identity, and developmental processes (Lieberman-Aiden et al., 2009; Dixon et al., 2012; Rao et al., 2014; Deng et al., 2012; Dixon et al., 2015; Bonev et al., 2017). Although chromatin architecture is dynamically regulated across biological contexts, inherited genetic variation represents an additional yet less understood source of architectural diversity (Grubert et al., 2020; Gorkin et al., 2019; C. Li et al., 2024; Spielmann et al., 2018; Sreenivasan et al., 2025). Increasing evidence links variation in chromatin architecture to inter-individual regulatory differences, disease pathogenesis, and evolutionary divergence (Kasowski et al., 2013; Lupiáñez et al., 2015; Javierre et al., 2016; Franke et al., 2016; Akdemir et al., 2020; Luo et al., 2021; Zemke et al., 2023; Che et al., 2026). Yet most chromatin interaction maps are interpreted within a standard singular reference coordinate system, leaving unresolved how naturally occurring genetic variation shapes 3D genome organization in the native context of individual haplotypes.

Structural variants (SVs) are among the largest and most consequential forms of human genetic variation, altering sequence content, dosage, spacing, and regulatory context (Alkan et al., 2011; Weischenfeldt et al., 2013; Sudmant et al., 2015; Chiang et al., 2017; Collins et al., 2020; Lupiáñez et al., 2015). Individual examples have shown that SVs can disrupt chromatin contacts, rewire regulatory landscapes, and alter transcriptional programs (Gröschel et al., 2014; Franke et al., 2016; Hnisz et al., 2016; Dixon et al., 2018; Guo et al., 2015). However, these examples do not establish how frequently inherited SVs influence chromatin organization across human populations. Because conventional chromatin analyses typically project structurally diverse haplotypes onto a standard reference genome, they may preserve broad interaction patterns while obscuring localized haplotype-specific chromatin differences (Ay & Noble, 2015; Zhang et al., 2024; Corda et al., 2025; Liu et al., 2026). Consequently, it remains unclear whether SVs broadly perturb 3D genome organization or whether only specific classes of variants, within particular genomic and regulatory contexts, contribute to recurrent chromatin contact divergence.

Recent advances in long-read sequencing, telomere-to-telomere (T2T) genome assembly, and haplotype-resolved genomics have transformed the ability to characterize human SVs at population scale (Audano et al., 2019; Ebert et al., 2021; Nurk et al., 2022; Liao et al., 2023; Logsdon et al., 2025; Lucas et al., 2026; Lin et al., 2026). In parallel, chromatin conformation profiling now enables genome architecture to be interrogated in genetically resolved contexts (Du et al., 2017; Richer et al., 2023; Tan et al., 2018; Liu et al., 2026). Together, these developments create an opportunity to move beyond reference-centered chromatin maps and ask how inherited SVs shape 3D genome organization across diverse human haplotypes.

Here, we use donor-specific assemblies (DSAs) generated by the Human Pangenome Reference Consortium (HPRC) to analyze haplotype-resolved chromatin interaction maps in their native genomic contexts at population scale and integrate these data with matched SVs, long-read transcriptomes, and comparative ape chromatin maps (Nurk et al., 2022; Liao et al., 2023; Yoo et al., 2025; Lucas et al., 2026; Lin et al., 2026). We examine how reference choice affects contact characterization and inferred 3D genomic structure, whether haplotype-specific chromatin contact differences are pervasive or concentrated into recurrent hotspots, and which SVs are associated with them. We then incorporate long-read transcriptomes from matched individuals to assess associations with gene-expression variation and comparative ape data to infer the evolutionary origins of associated SV alleles. Together, this study establishes a haplotype-resolved framework for distinguishing widespread sequence variation from the restricted subset of SVs associated with recurrent chromatin contact divergence and expression variation across human populations and evolution.

## RESULTS

### Haplotype-resolved mapping reveals reference-dependent distortion of chromatin contacts

To determine how structural diversity of genomes affects the inference of 3D genome organization, we generated deeply sequenced Hi-C/Omni-C data from lymphoblastoid cell lines (LCLs) of 177 genetically diverse individuals whose genomes were assembled as part of the HPRC Release 2 (Figure 1A) (Lucas et al., 2026). These data comprise 148.6 billion raw read pairs, with an average of 839.5 million read pairs per individual (Table S1). The datasets are of high quality, with strong replicate concordance (median GenomeDISCO reproducibility score, 0.911; interquartile range (IQR), 0.904–0.919), a high cis-interaction fraction (mean 74%, range 53–87%), and high contact-map resolution (median 3.95 kbp) (Figures S1A–S1C; Table S2). Each individual is represented by two phased haplotype assemblies, enabling chromatin contacts to be compared after alignment to a shared reference genome or to the corresponding donor-specific haplotypes. For trio-phased samples, Hap1 and Hap2 correspond to the paternal and maternal haplotypes, respectively; for non-trio samples, the autosomal Hap1/Hap2 assignment is arbitrary, with the exception that the Y chromosome (chrY) is by convention assigned to Hap1 in males. Throughout, Hap1 and Hap2 therefore refer to within-individual phasing labels and do not denote particular parental origin across the majority of samples.

**Figure 1.**
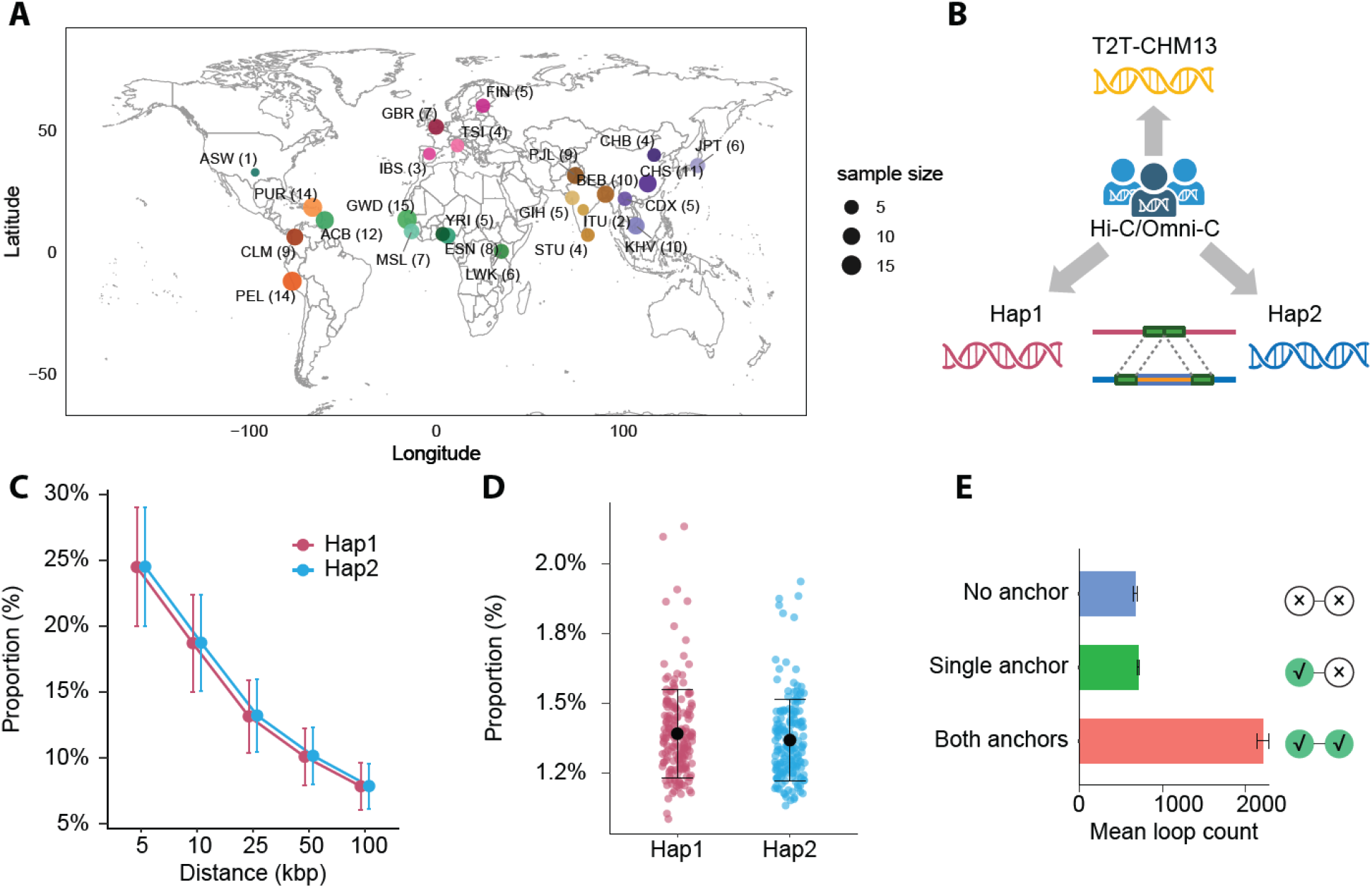
Haplotype-resolved mapping reveals reference-dependent shifts in chromatin contacts. (A) Geographic and population distribution of the 177 HPRC Release 2 individuals analyzed in this study. HG002 is not shown because geographic metadata of the collection site is unavailable. Colors distinguish populations; point size is scaled by sample size, with the exact number of individuals in each population shown in parentheses. Population abbreviations follow the 1000 Genomes Project nomenclature (Auton et al., 2015). (B) Schematic of the mapping strategy. Chromatin interaction reads from each individual were aligned to the shared T2T-CHM13 reference and to the two corresponding DSAs. (C) Proportion of shared cis Hi-C contact pairs for which the inferred genomic distance differs as function of kbp between DSA mapping and T2T-CHM13 mapping by at least the indicated threshold. For each shared cis contact pair, genomic contact distance was calculated as the absolute distance between the mapped positions of the two interaction anchors. Contact-distance shift is defined as the absolute difference between the inferred contact distances in the DSA and T2T-CHM13 coordinate systems. Points and error bars show the mean ± standard deviation (s.d.) across individuals, calculated separately for Hap1 and Hap2. (D) Proportion of valid contact pairs recovered by DSA mapping only but not by T2T-CHM13 mapping for Hap1 and Hap2. Colored points represent individual haplotypes, shown separately for Hap1 and Hap2; black points and error bars indicate the mean ± s.d. (E) Loop-anchor sharing between Hap1 and Hap2. Loop anchors are defined as the genomic intervals at the two ends of each called loop. Loops are classified according to whether neither, one, or both anchors overlapped anchors in the other haplotype’s loop set. Hap1 and Hap2 loop counts are averaged within each individual. Bars show the mean across individuals, and error bars indicate ±1 standard error of the mean (s.e.m.). Schematics indicate matched (check marks) and unmatched (crosses) anchors. HPRC, Human Pangenome Reference Consortium; T2T-CHM13, telomere-to-telomere CHM13 reference genome; DSA, donor-specific assembly; Hap1 and Hap2, haplotypes 1 and 2, respectively.

We mapped each dataset to the T2T-CHM13 reference and to the two corresponding DSAs to compare contact pairs recovered across coordinate systems (Figure 1B; Table S3). Most valid pairs are shared between DSA-based and T2T-CHM13-based mappings, with an average of 98.6% (median, 98.6%; IQR, 98.5–98.7%) of DSA-valid pairs also recovered after T2T-CHM13 mapping, indicating that broad chromatin interaction patterns are largely preserved irrespective of reference (Figures S2A). However, shared cis contacts frequently show substantial differences in inferred genomic distance, with approximately 19% differing by more than 10 kbp (median across haplotypes: Hap1, 18.3%, IQR, 15.8–21.4%; Hap2, 18.4%, IQR, 16.2–21.3%) and approximately 8% differing by more than 100 kbp (Hap1, 7.7%, IQR, 6.5–9.0%; Hap2, 7.9%, IQR, 6.5–9.2%) between mapping contexts (Figures 1C and S2B; Table S4). Thus, a shared reference preserves most contact identities yet substantially misestimates genomic separation for a sizable subset of shared cis contacts.

DSA-based mapping also recovers an additional 2.8% (average 1.39% in Hap1 and 1.37% in Hap2, compared to T2T-CHM13) of valid contact pairs that are not retained after alignment to T2T-CHM13 (Figure 1D). These DSA-specific pairs represent candidate contacts involving haplotype-specific or highly divergent sequence contexts that are incompletely captured by a single standard reference. Although these contacts constitute a minority of all valid pairs, their recovery indicates that reference-based analyses may miss nearly 3% localized interaction information.

We next asked whether reference-dependent contact shifts affect inferred chromatin features. Across the 354 haplotypes, a median of 60.0% of loops called from DSA-based maps matched loops called from the corresponding T2T-CHM13-based map at both anchors, whereas 18.3% match only one anchor and 21.1% show no anchor-level match (Figure S2C). Similarly, reciprocal comparison of loop calls between the two haplotypes of each individual show that a median of 60.6% of loops match a loop on the alternate haplotype at both anchors, whereas 18.0% match at only one anchor and 20.9% at neither anchor (Figure 1E), consistent with localized haplotype-dependent differences in loop anchoring.

### Haplotype-specific chromatin contact divergence occurs in discrete hotspots

To determine how chromatin contacts differ between the two haplotypes of each individual, we first identified all haplotype-specific contact pairs across the genome (Figure 2A). Approximately 99% of contacts are shared between haplotypes, whereas a minority of contacts (median Hap1: 1.02%, Hap2: 1.11%) are detected specifically in one haplotype (Figure S3A). These haplotype-specific contacts are largely supported by uniquely mappable sequences, with approximately 99.9% of contact ends and approximately 99.8% of complete contact pairs falling within uniquely mappable regions (Figure S3B). Remapping haplotype-specific reads to the alternate haplotype assembly further showed that 98.7% did not align concordantly: 9.7% were unmapped and 89.0% had only one read end aligned (Figure S3C), arguing against cross-haplotype mapping ambiguity as the predominant source of haplotype-specific contacts.

**Figure 2.**
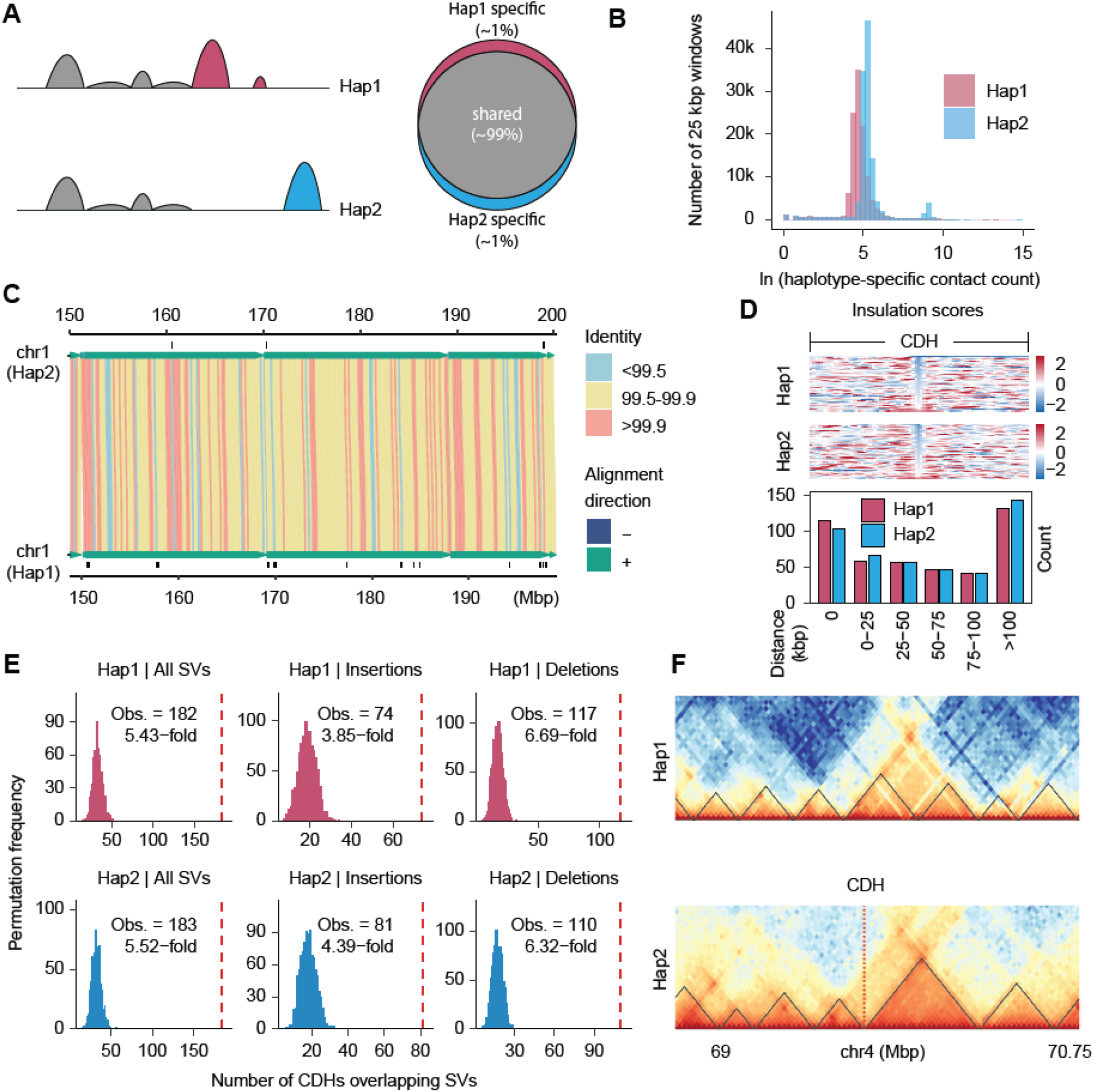
Haplotype-specific chromatin contact divergence occurs at discrete hotspots. (A) Schematic of shared and haplotype-specific chromatin contacts between the two phased haplotypes of an individual. For each haplotype, approximately 99% of contacts are shared with the other haplotype, whereas approximately 1% are haplotype-specific. (B) Distribution of haplotype-specific contact counts in 25 kbp genomic windows for Hap1 and Hap2. The x-axis shows the natural logarithm of the haplotype-specific contact count per 25 kbp window. The secondary peak in Hap2 reflects elevated haplotype-specific contact counts in X chromosome windows. (C) Distribution of CDHs across chr1: 150–200 Mbp in a representative individual (HG002). Black tick marks indicate CDH locations. Sequence identity and alignment orientation between the two haplotypes across the interval are shown. (D) Relationship between CDHs and TAD boundaries. Top, insulation-score profiles in Hap1 and Hap2. Each row represents a CDH located within 25 kbp of its nearest TAD boundary and is aligned to the boundary midpoint (±1 Mbp). Rows are ordered by the score at the central position. The color scale represents insulation scores. Bottom, distribution of distances from CDHs to the nearest TAD boundary. (E) Permutation analysis of overlap between CDHs and heterozygous SVs in HG002. Columns show insertions and deletions combined (left), insertions (middle), and deletions (right). Top, Hap1; bottom, Hap2. Histograms show null distributions from 1,000 chromosome- and length-preserving permutations of CDH windows. Dashed red lines indicate the observed numbers of CDHs overlapping at least one SV. (F) Representative locus on chromosome 4 showing haplotype-specific chromatin contact divergence near a CDH. A CDH detected in Hap2 overlaps a TAD boundary (chr4_RagTag: 69,712,500–69,737,500), whereas the corresponding region in Hap1 lacks the CDH. The heterozygous sequence is absent from Hap1 and is represented as an insertion in Hap2 relative to the T2T-CHM13 reference (chr4-69535832-INS-60). The Hap2 CDH is marked by the dashed red line. CDH, contact difference hotspot; TAD, topologically associating domain; SV, structural variant; Obs., observed number of CDHs overlapping at least one SV.

We next asked whether haplotype-specific contacts were distributed uniformly across the genome or concentrated within discrete regions. Genome-wide quantification of haplotype-specific contact counts in 25 kbp windows reveals a unimodal distribution with a pronounced right tail, consistent with spatial heterogeneity in local contact enrichment (Figure 2B). We, therefore, developed a negative-binomial framework to identify genomic regions significantly enriched for haplotype-specific contacts (Methods). To minimize confounding by mapping and assembly artifacts, we excluded low-coverage windows, sex chromosomes and acrocentric regions (short arms of chromosomes 13, 14, 15, 21, and 22). In total, we identified a median of 580 contact difference hotspots (CDHs) per haplotype (IQR, 526–631), which merged into a median of 438 contiguous regions (IQR, 401–467; Figure 2C; Table S5).

CDHs are highly reproducible across contact-distance and coverage filters, with a median of 86% of CDHs called at 25 kbp resolution also recovered at 10 kbp resolution (range, 76–94%; Figures S4A–S4C). CDHs show little evidence of preferential localization to sequence contexts prone to mapping artifacts, with only modest enrichment in segmental duplications (SDs) (median, 1.31-fold; range, 0.91–1.64), overall depletion from satellite repeats (median, 0.85-fold; range, 0.52– 1.03), and minimal enrichment across dispersed repeat classes (median enrichments, 1.02–1.05-fold across classes) (Figures S4D–S4F). These results argue against technical artifacts as the predominant source of CDHs.

We then examined whether CDHs coincide with annotated 3D genome features. Under chromosome-matched permutation, CDHs show a median 1.28-fold enrichment at topologically associating domain (TAD) boundaries and a 1.28-fold enrichment at loop anchors relative to randomized regions, with a median of 15.2% and 12.7% of CDH regions directly overlapping TAD boundaries and loop anchors, respectively (Figure 2D). These positional associations indicate that CDHs co-localize with features of local genome organization.

We further compared the distribution of heterozygous SVs with the map of CDHs. Heterozygous insertions and deletions at least 50 bp were identified using a comprehensive combination of read- and assembly-based approaches as part of the HPRC variant discovery framework (Methods) (Lin et al., 2026). Relative to matched genomic permutations, heterozygous deletions are enriched 8.3-fold within CDHs and heterozygous insertions 5.3-fold (empirical *P* < 0.001 for both; Figure 2E; Table S6). This enrichment is consistently observed across haplotypes (Figure S5A). Nevertheless, only 2.9% of heterozygous insertions and 4.1% of heterozygous deletions intersect CDHs, indicating that most heterozygous SVs do not coincide with detectable haplotype-specific contact enrichment (Figure S5B). Thus, chromatin contact divergence is associated with a selective subset of SV-containing genomic contexts rather than reflecting a general property of all heterozygous SVs.

A representative locus in HG002 illustrates this pattern. Although chromatin contacts are broadly shared between the two haplotypes, a CDH is detected in only Hap2 and coincides with a TAD boundary (boundary position: chr4_RagTag: 69712500-69737500) (Figure 2F). Hap2 carries a 60 bp insertion at this locus (T2T-CHM13 variant ID: chr4-69535832-INS-60), whereas the corresponding region in Hap1 lacks both the insertion and a detected CDH. These observations illustrate a locus-specific association between a heterozygous SV and localized haplotype-specific contact enrichment.

### Chromatin-structure–associated SVs mark recurrent CDHs

To determine whether CDHs represent private haplotype-specific events or recurrent features of population variation, we performed a pangenome analysis across all 354 haplotypes. More than 99% of CDHs could be projected from DSAs onto T2T-CHM13, enabling comparison in a common genomic coordinate system (Figure S6A). Clustering overlapping CDHs across haplotypes identified 1,790 recurrent CDH regions, substantially more than expected under matched genomic permutations (1,790 observed versus 225 ± 75 permuted regions, mean ± s.d.; empirical *P* = 0.001, 999 permutations; Figures 3A and S6B; Table S7). Thus, haplotype-specific chromatin contact divergence repeatedly converges on a restricted set of genomic loci across the population.

**Figure 3.**
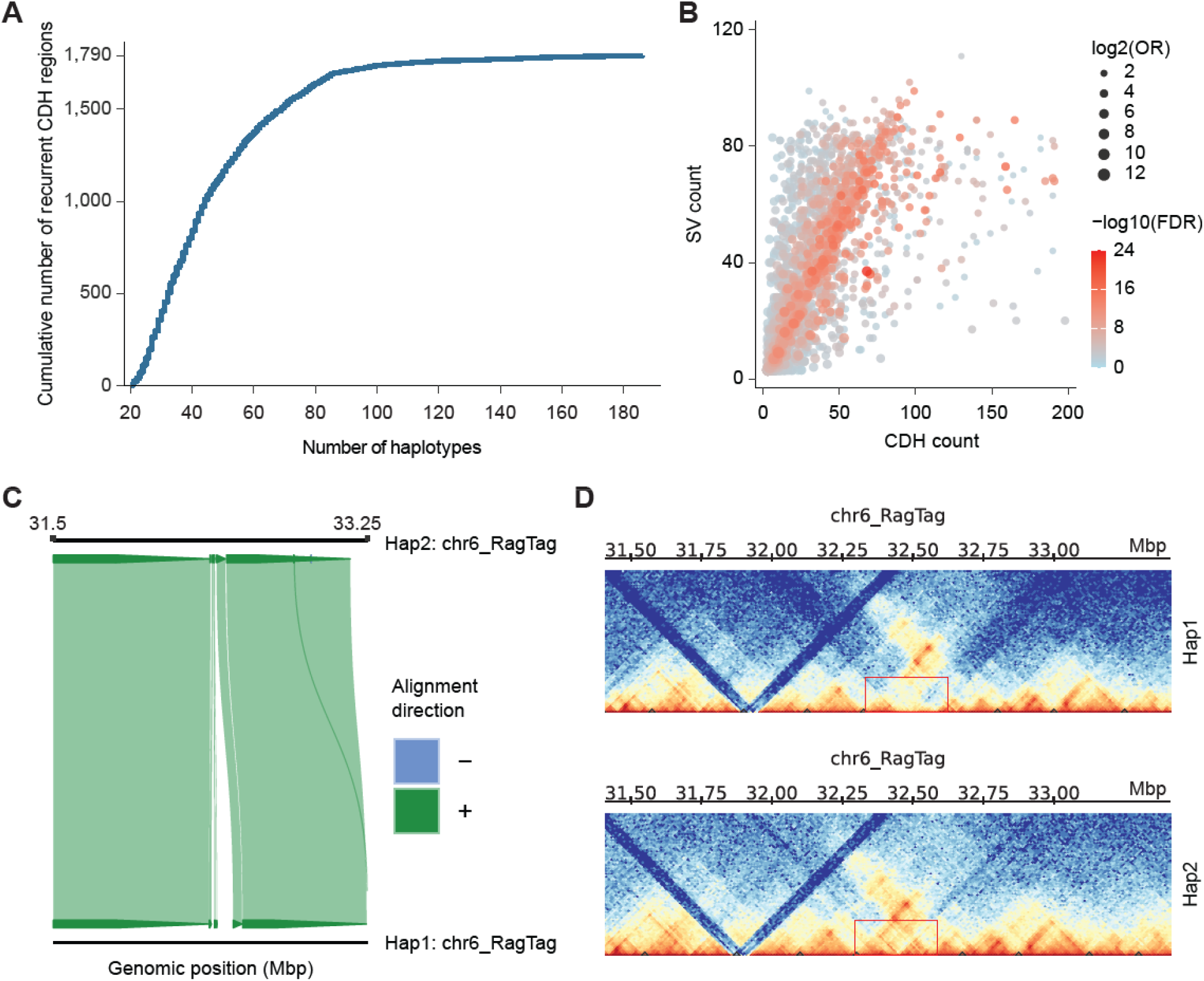
CSA-SVs mark recurrent CDHs. (A) Cumulative distribution of haplotype support across 1,790 recurrent CDH regions in T2T-CHM13 coordinates. For each region, haplotype support was defined as the largest number of haplotypes whose CDHs overlap at the same position. The curve shows the cumulative number of regions with support value shown on the x-axis. (B) Association between SV occurrence and CDHs across haplotypes. Each point represents an SV locus; the x-axis shows the number of haplotypes with a CDH at the corresponding interval, and the y-axis shows the number of haplotypes carrying the SV. Point size indicates log2 (OR), and color indicates −log10 (FDR). (C) Haplotype assembly alignment across the region surrounding a representative 1.6 kbp insertion at chr6:32,378,735 in HG002. The insertion is present in Hap2 but absent from Hap1. Alignment direction is indicated by color. (D) Haplotype-resolved chromatin contact maps across the corresponding region in HG002. In Hap2, the insertion coincides with a CDH and pronounced enrichment of haplotype-specific contacts across the surrounding region, whereas Hap1 lacks both the insertion and the corresponding CDH and shows substantially lower local contact enrichment. Red boxes mark the region of haplotype-specific chromatin contact divergence. CSA-SV, chromatin-structure–associated structural variant; OR, odds ratio; FDR, false discovery rate.

We next integrated the pangenome CDH map with the HPRC SV callset to identify SVs associated with recurrent chromatin contact divergence (Lin et al., 2026). For each nonredundant SV locus, we tested whether haplotypes carrying the variant were more likely to contain a CDH at the corresponding genomic interval than haplotypes lacking the variant (Methods). This analysis identified 2,495 nonredundant SV loci, including 810 insertions and 1,685 deletions, whose presence across haplotypes was significantly associated with CDH occurrence (odds ratio (OR) > 2 and false discovery rate (FDR)-adjusted *P* < 0.05; two-sided Fisher’s exact test with Benjamini–Hochberg correction; Figure 3B; Table S8). We refer to these candidate loci as chromatin-structure–associated SVs (CSA-SVs).

Across the 354 haplotypes, each haplotype carries a median of 270 CSA-SVs (IQR, 249–290) (Figure S7A). These CSA-SVs range from 50 bp to 70.3 kbp, with a median size of 2.11 kbp (IQR, 0.74–4.29 kbp). Adjustment for local genetic ancestry had little impact on CSA-SV effect estimates, which were highly correlated before and after adjustment (r = 0.986), with 99.4% retaining the same direction (Figure S7B). In a complementary within-individual analysis, treating the two haplotypes from each individual as correlated increased standard errors only modestly (median, 1.15-fold), with all associations remaining significant, indicating that within-individual haplotypes had little impact on the observed associations (Figure S7C). These analyses support the robustness of CSA-SV associations to adjustment for local ancestry and within-individual dependence.

To characterize the local chromatin context of CSA-SVs, we quantified haplotype-specific contact density around each SV locus. The median contact density around CSA-SVs is 6.8-fold higher than around CDH-overlapping SVs that do not meet the CSA-SV criteria (two-sided Wilcoxon signed-rank test, *P* < 0.001), with the same trend observed for both insertions and deletions (Figure S7D). In addition, a median of 28.4% of CSA-SVs overlap or occur within 25 kbp of chromatin architectural features, including TAD boundaries and loop anchors (Figure S7E). Thus, CSA-SVs are characterized by pronounced local enrichment of haplotype-specific contacts and frequently localized near established features of chromatin organization. Most CSA-SVs map outside protein-coding exons (99%), placing them predominantly in noncoding regions (Figure S7F).

A representative locus on chromosome 6 illustrates the relationship between a heterozygous SV and local chromatin contact divergence (Figure 3C). In Hap2, a 1.6 kbp insertion at chr6:32,378,735 coincides with a CDH and pronounced enrichment of haplotype-specific contacts across the surrounding region. By contrast, Hap1 lacks both the insertion and the corresponding CDH and shows substantially lower local contact enrichment (Figure 3D). This haplotype-level correspondence is consistent with a localized association between the insertion and chromatin contact divergence.

### CSA-SVs link CDHs to gene-expression variation

To determine whether CSA-SVs are associated with gene-expression variation, we integrated long-read RNA-sequencing (RNA-seq) data from a matched subset of 163 individuals used in the haplotype-resolved chromatin analyses (Table S9). This dataset consists of 2.87 billion full-length non-chimeric (FLNC) reads, enabling population-scale analysis of gene expression variation in relation to CSA-SV genotypes.

Cis-association analysis identified 237 genotype–expression associations, encompassing 195 genes whose expression differed significantly among CSA-SV genotypes (Kruskal–Wallis test, Benjamini–Hochberg FDR < 0.05; Table S10). Expression differences associated with CSA-SVs are frequently dosage-dependent, with 175 of 237 significant associations also showing a significant dosage trend (Benjamini–Hochberg FDR < 0.05; Figure 4A). Among associations for which all three genotype classes are represented and median expression levels are distinct, 83.3% showed a monotonic relationship between alternative-allele dosage and median expression, substantially exceeding the 33.3% expected under random ordering (binomial test, *P* = 1.9 × 10 ²; Figure S8A and S8B). At representative loci such as *SESTD1* and *SIGLEC5*, expression levels differ among CSA-SV genotype classes (Figure 4B), illustrating locus-specific relationships between CSA-SV genotype and gene-expression variation.

**Figure 4.**
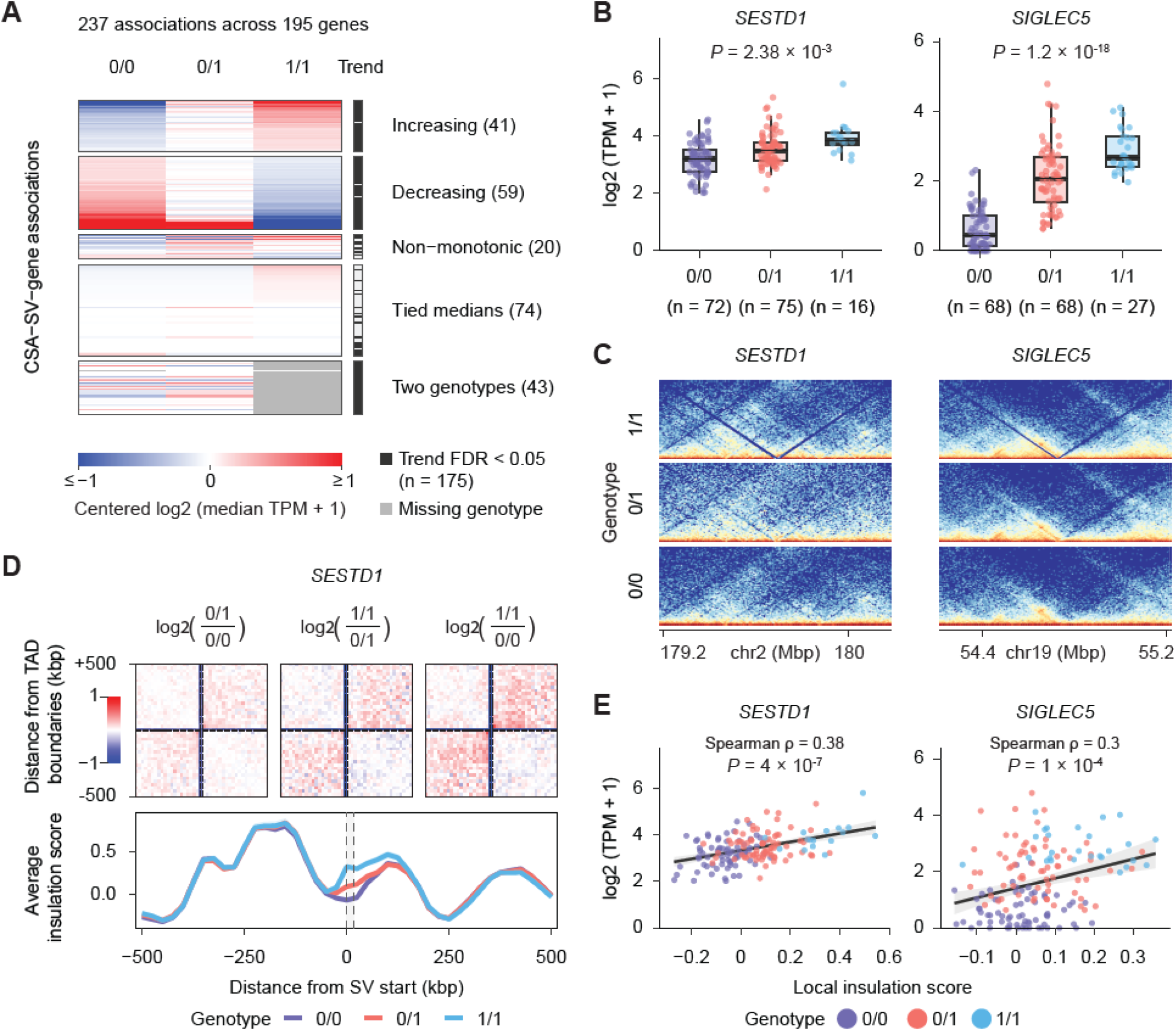
CSA-SVs associate with expression differences. (A) Genotype-associated expression patterns for 237 CSA-SV–gene associations encompassing 195 genes (Kruskal–Wallis test, Benjamini–Hochberg FDR < 0.05). Rows represent associations, and columns represent CSA-SV genotype classes (0/0, 0/1, and 1/1). Colors indicate log (median TPM + 1), centered within each row across observed genotype classes, with the color scale saturated at −1 and 1. Associations are grouped into increasing, decreasing, non-monotonic, tied-median, and two-genotype categories. Gray heatmap cells indicate missing genotype classes. Dark gray marks in the adjacent annotation strip indicate significant dosage trends (Benjamini–Hochberg FDR < 0.05). (B) Expression of *SESTD1* and *SIGLEC5* across CSA-SV genotype classes. These expression patterns show cis-association with context-dependent SVs (*SESTD1*, chr2-179683834-DEL-16208; *SIGLEC5*, chr19-54718506-DEL-16422). Boxes show the median and IQR; whiskers indicate 1.5× the IQR. (C) Genotype-stratified chromatin contact maps surrounding the *SESTD1*-associated (left) and *SIGLEC5*-associated (right) CSA-SVs. Each map shows the CSA-SV and 500 kbp of flanking sequence on either side. Rows show genotype classes 1/1, 0/1, and 0/0 from top to bottom. Colors indicate contact intensity. (D) Genotype-associated chromatin organization at the *SESTD1* locus. Top, log ratios of aggregate chromatin contact maps comparing 0/1 with 0/0, 1/1 with 0/1, and 1/1 with 0/0 (left to right). Red and blue indicate higher and lower contact frequencies, respectively, in the numerator genotype. Bottom, mean insulation-score profiles for the three genotype classes. Positions are shown relative to the SV start. (E) Relationship between local insulation score and expression of *SESTD1* and *SIGLEC5*. Each point represents one individual, colored by CSA-SV genotype. Expression is shown as log (TPM + 1), and local insulation scores are taken from the genomic bin containing the SV start. Black lines show linear fits across individuals, with gray bands indicating 95% confidence intervals. Spearman correlation coefficients and two-sided *P* values are shown. TPM, transcripts per million; IQR, interquartile range; 0/0, homozygous for the reference allele; 0/1, heterozygous for the reference and alternate alleles; 1/1, homozygous for the alternate allele.

Using population-scale chromatin contact maps, we performed aggregate insulation analyses which revealed genotype-dependent differences around CSA-SVs, with distinct local insulation profiles observed for non-carriers, heterozygous carriers, and homozygous carriers (pairwise Mann–Whitney U tests with Benjamini–Hochberg correction, FDR < 0.001; Figures 4C, 4D, and S9A–S9C). These findings link CSA-SV genotypes to local variation in domain insulation, providing a potential structural correlate of the expression differences observed at these loci.

Finally, we evaluated the relationship between local chromatin architecture and gene expression by correlating insulation scores with gene-expression levels across individuals. For both the *SESTD1* (Spearman ρ = 0.385, *P* = 4 × 10 ^7^) and *SIGLEC5* loci (Spearman ρ = 0.299, *P* = 7.68 × 10), we observed significant correlations between local chromatin insulation and gene expression (Figure 4E).

### Human-derived CSA-SVs associate with regulatory and phenotypic variation

To investigate the evolutionary origins of CSA-SVs, we analyzed the orthologous loci across six nonhuman ape species—chimpanzee, bonobo, gorilla, Bornean orangutan, Sumatran orangutan and siamang—using haplotype-resolved T2T assemblies and matched chromatin contact maps (Methods) (Yoo et al., 2025). This comparative framework enabled us to examine both the inferred evolutionary origin of CSA-SV alleles and their association with human-lineage chromatin contact divergence. Among CSA-SVs with confidently assigned evolutionary origins, 93.0% (1,395/1,500) are inferred to be derived in the human lineage. Across the full CSA-SV set, ape-matching allelic states are detected for 6.5% of insertions and 3.1% of deletions (Figure 5A; Table S11). Human-derived CSA-SVs are significantly larger than ancestral CSA-SVs (median, 2.61 kbp versus 0.33 kbp; two-sided Wilcoxon rank-sum test, *P* = 1.5 × 10 ²; Figures 5B, S10A, and S10B). These patterns suggest that CSA-SVs are predominantly human-lineage derived and tend to involve larger genomic segments than their ancestral counterparts.

**Figure 5.**
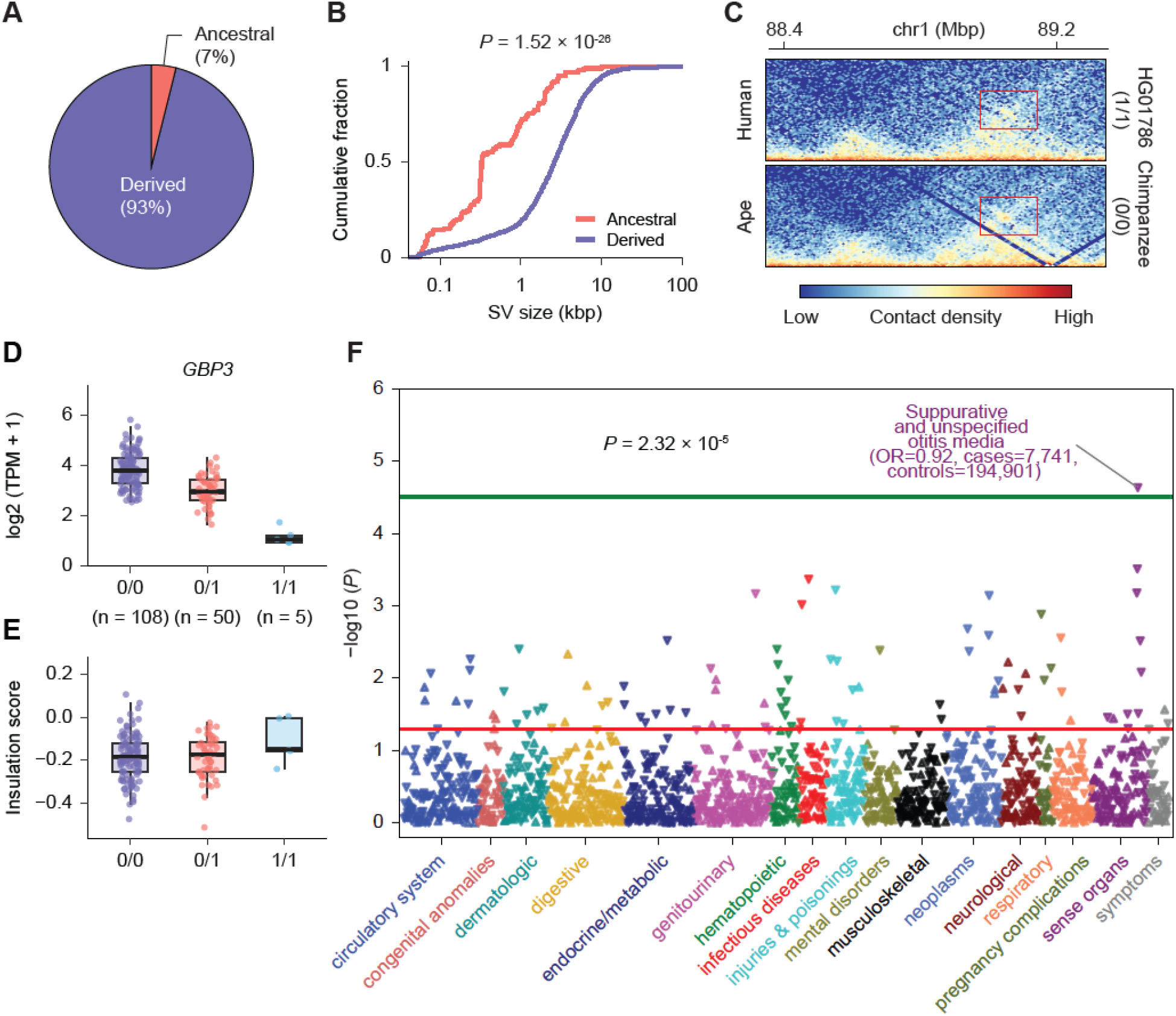
Human-derived CSA-SVs link to local chromatin and regulatory divergence. (A) Inferred ancestral and human-derived states of CSA-SV deletions and insertions based on orthologous ape allelic states. (B) Empirical cumulative distributions of SV sizes for ancestral (coral; n = 105) and human-derived (purple; n = 1,395) CSA-SVs with confidently assigned evolutionary origins, pooling insertions and deletions. Curves show the fraction of CSA-SVs with sizes at or below each value. SV sizes are shown on a logarithmic scale. *P* = 1.52 × 10 ², two-sided Wilcoxon rank-sum test. (C) Representative human–chimpanzee chromatin contact maps surrounding a 2.7 kbp human-derived deletion at chr1:88,853,624–88,856,340 (T2T-CHM13). Maps are displayed at 10 kbp resolution and span 500 kbp on either side of the deletion start. Both maps are shown in T2T-CHM13 coordinates. Red boxes highlight local contact differences corresponding for comparison, and colors indicate contact density. This deletion is associated with expression variation in *GBP3*, whose expression levels across deletion genotypes are shown in (D). (D) *GBP3* expression across genotypes of the 2.7 kbp human-specific deletion at chr1:88,853,624–88,856,340. Boxes show the median and IQR; whiskers indicate 1.5× the IQR. (E) Insulation scores in the window containing the deletion start at the *GBP3* locus, stratified by deletion genotype (0/0, 0/1, and 1/1). Points represent individual samples (n = 108, 50 and 5, respectively). Boxes show the median and IQR; whiskers extend to the most extreme observations within 1.5× the IQR of the box limits. (F) Phenome-wide association analysis of the *GBP3*-associated deletion in the *All of Us* cohort. The green line indicates the multiple-testing significance threshold.

We next asked whether human-derived CSA-SVs are associated with chromatin contact differences across species. At a locus harboring a 2.7 kbp human-derived deletion that remains polymorphic in modern humans, contact maps reveal localized differences between a human individual homozygous for the derived deletion and a chimpanzee retaining the inferred ancestral, undeleted allele (Figure 5C). These observations illustrate that human-derived CSA-SVs can coincide with local chromatin contact divergence between humans and nonhuman apes.

This human-derived CSA-SV is associated with expression variation of *GBP3*, which encodes an interferon-inducible GTPase involved in innate immune defense (Nordmann et al., 2012). Among individuals with matched RNA-seq data, homozygous carriers of the ancestral allele show the highest *GBP3* expression, whereas heterozygous and homozygous deletion carriers show progressively reduced expression levels (Figure 5D). This deletion, thus, corresponds to a significant cis association with *GBP3* expression (*P* < 0.001) as well as genotype-dependent differences in local chromatin contacts (Figure 5E). It is interesting that our phenome-wide association analysis (PheWAS) of the deletion allele in the *All of Us* cohort suggests a reduced risk of suppurative and unspecified otitis media after multiple-testing correction, consistent with this gene’s function in immunity (OR = 0.92; *P* = 2.32 × 10; 7,741 cases and 194,901 controls; Figure 5F). While this association does not establish causality, it provides population-level support for a link between a human-derived CSA-SV and phenotypic variation.

## DISCUSSION

This study provides insight into a long-standing question of whether and to what extent inherited SVs reshape chromatin organization across human populations. Our analyses suggest that SVs influence chromatin organization in a selective rather than pervasive manner, with only a restricted subset of variants associated with recurrent chromatin contact divergence. The identification of CSA-SVs, which show association with chromatin contact differences in the population, supports a model in which chromatin contact divergence is concentrated within discrete genomic contexts and reflects selective interactions between SV and regulatory architecture. This interpretation is consistent with experimental evidence that the regulatory consequences of SVs depend on local chromatin architecture and enhancer–promoter context (Franke et al., 2016; Despang et al., 2019; Xu et al., 2022). Consistent with this view, CSA-SVs are linked to expression variation and exhibit evolutionary signatures, supporting a broader role for SVs in shaping regulatory diversity.

Our findings suggest that analyses limited to a standard reference genome may underestimate the full extent of chromatin contact divergence across individuals. Unlike measurements interpreted locally along the linear genome, chromatin interaction maps highly depend on the accurate coordinate placement of distant genomic loci and may therefore be particularly sensitive to genomic structural differences between haplotypes (Liu et al., 2026; Macias-Velasco et al., 2026). As a consequence, analyses performed within a single reference coordinate system may preserve broad interaction patterns while obscuring localized haplotype-specific differences in chromatin organization. By incorporating haplotype-resolved genome assemblies, our study extends the current framework for chromatin interpretation and suggests that personalized DSA references can improve the precision of characterizing chromatin variation across populations.

More broadly, our findings establish a population-scale map of SV-associated 3D genome variation. As the CSA-SVs show potential functional effects, the multi-omics approach combining haplotype-resolved genome assemblies, chromatin contact maps, and transcriptomes from the same donor enables the prioritization of SVs of functional consequence. While a typical human genome has ∼25,000 SVs, most of these are predicted to be neutral and the approach outlined here may be a more general strategy to hone in on SVs that link gene expression (Audano et al., 2019; Ebert et al., 2021; Scott et al., 2021; Logsdon et al., 2025). The *GBP3* locus further illustrates how this framework can identify variants associated with both gene-expression and phenotypic variation, while leaving the causal relationships among these observations to be resolved.

This study has several limitations. Our primary analyses were restricted to LCLs and to the limited set of nonhuman ape samples for which matched haplotype-resolved assemblies and chromatin maps are currently available. Moreover, the observed associations among SVs, chromatin contact divergence, and gene-expression differences do not establish causality for individual variants but instead identify candidates for functional follow-up. An important next step will be to extend this framework to the Somatic Mosaicism across Human Tissues (SMaHT) resource, which is generating genomic, chromatin, and transcriptional profiles across multiple normal tissues from the same donors (Coorens et al., 2025). These analyses could assess the tissue specificity and reproducibility of candidate relationships, whereas targeted perturbation of prioritized loci in relevant cell types will still be required to demonstrate causality. Finally, extending this strategy to larger and more diverse human cohorts, together with additional comparative primate datasets, will clarify the generality and evolutionary relevance of SV-associated chromatin variation.

## METHODS

### Sample selection

#### Human samples

Haplotype-resolved genome assemblies were obtained from HPRC Release 2 (Lucas et al., 2026). Individuals were included if matched Omni-C/Hi-C data from LCLs were available, yielding 177 individuals represented by 354 phased haplotype assemblies. Sample identifiers, population information, sequencing yields, library types, and replicate counts are provided in Table S1.

#### Ape samples

Haplotype-resolved genome assemblies and matched Omni-C data were obtained from the ape T2T genome project. The dataset comprised six individuals, one from each of six species: chimpanzee, bonobo, gorilla, Bornean orangutan, Sumatran orangutan, and siamang. Each individual was represented by two phased haplotype assemblies, providing 12 nonhuman ape haplotypes for comparative analyses. Sample information, sequencing assays, and data accession numbers are described in the original study (Yoo et al., 2025).

### Chromosome conformation capture

Omni-C libraries were prepared using the Dovetail Omni-C kit (Dovetail Genomics) with minor modifications. Briefly, 1 × 10 cells were fixed with formaldehyde and DSG, and chromatin was digested with DNase I to generate appropriately sized fragments. Chromatin ends were repaired, ligated to a biotinylated bridge adaptor, and proximity ligated. Crosslinks were then reversed, and proximity-ligated DNA was purified. Illumina sequencing libraries were prepared using the NEBNext Ultra II library preparation kit (NEB) with Y-adaptors. Biotin-containing ligation products were enriched by streptavidin bead capture. Each capture reaction was split into two technical library replicates before final PCR enrichment to preserve library complexity. Libraries were uniquely dual indexed and sequenced on an Illumina NovaSeq platform with 2 × 150-bp paired-end reads.

### Preparation of DSA mapping references

Contigs from each phased haplotype assembly were scaffolded independently using RagTag v2.1.0 (Alonge et al., 2022), with T2T-CHM13 v2.0 as the guide reference. Scaffolding was performed using ragtag.py scaffold with default parameters. The resulting scaffolded assemblies were used as DSA mapping references.

### Omni-C/Hi-C data processing

Reads were processed using a workflow adapted from the Dovetail Omni-C analysis pipeline (https://dovetail-analysis.readthedocs.io/en/latest/). For each individual, read pairs were independently aligned to T2T-CHM13 v2.0 and to each of the two corresponding DSAs using BWA-MEM v0.7.17 with the parameters −5SP -T0 (H. Li, 2013). Alignments were processed using pairtools v1.0.3 (Open2C et al., 2024). Contact pairs were extracted using pairtools parse --min-mapq 40 --walks-policy 5unique --max-inter-align-gap 30, sorted using pairtools sort, and deduplicated using pairtools dedup --mark-dups with default duplicate-detection settings, including a maximum coordinate mismatch of 3 bp. Nonduplicate pairs with both reported ends uniquely mapped (MAPQ ≥40) were exported in .pairs format using pairtools split, and processing statistics were recorded using --output-stats. Duplicate pairs were additionally exported in a separate processing run using --output-dups --keep-parent-id, retaining the identifier of the representative read pair associated with each duplicate.

Contact-pair files were bgzip-compressed and indexed using Pairix v0.3.8 (Lee et al., 2022). Contact matrices were generated at 10, 25, and 100 kbp resolution using cooler cload pairix (cooler v0.9.3) (Abdennur & Mirny, 2020). Matrix-balancing weights were calculated using cooler balance. Raw matrices were also converted to HiCExplorer .h5 format using hicConvertFormat and normalized using the Knight–Ruiz algorithm with hicCorrectMatrix correct --correctionMethod KR (HiCExplorer v3.7.3) (Ramírez et al., 2018; Wolff et al., 2018, 2020). Multiresolution .hic files were generated using the pre command in Juicer Tools v1.22.01 (Durand et al., 2016).

### Data quality control

Reproducibility between replicates was assessed using GenomeDISCO with default parameters on 100 kbp contact maps generated from T2T-CHM13 mappings (Ursu et al., 2018). All distinct pairs of replicate datasets from the same individual were compared.

For contact-map resolution estimation, contact pairs retained after deduplication of each sequencing dataset were pooled for each individual. Resolution was estimated using the procedure described previously, which defines contact-map resolution as the smallest bin size at which at least 80% of genomic bins contain at least 1,000 contacts (Rao et al., 2014).

Cis- and trans-contact counts were obtained from pairtools summary statistics and summed across sequencing datasets for each individual. The cis-contact fraction was calculated as the number of nonduplicate intrachromosomal contact pairs divided by the total number of nonduplicate mapped contact pairs, including both cis and trans contacts.

### Identification of TAD boundaries

TAD boundaries were identified from 25 kbp contact matrices using hicFindTADs in HiCExplorer v3.7.3. TAD-separation scores were computed from z-score-transformed contact matrices across multiple window sizes. Candidate boundaries were identified as local minima in the mean TAD-separation score across window sizes and filtered using an FDR threshold of 0.05 and a delta threshold of 0.01. The following parameters were used: --minDepth 75000 --maxDepth 250000 --step 25000 --thresholdComparisons 0.05 -- correctForMultipleTesting fdr --delta 0.01. All other analysis parameters, including -- minBoundaryDistance, were left at their defaults. TADs were defined as genomic intervals between consecutive retained boundaries on the same chromosome.

### Identification of chromatin loops

Chromatin loops were identified separately for each individual from 10 kbp contact matrices using hicDetectLoops in HiCExplorer v3.7.3. The algorithm computes observed-to-expected contact values, preselects candidate interactions using distance-specific continuous negative-binomial models, and evaluates candidate peaks against local background regions using Wilcoxon rank-sum tests. Loop calling was restricted to intrachromosomal interactions within 2 Mbp using the following parameters: -- maxLoopDistance 2000000 --windowSize 10 --peakWidth 6 --pValuePreselection 0.05 –pValue 0.05. The windowSize and peakWidth parameters were specified in matrix bins, and all other analysis parameters were left at their defaults. The resulting loop calls included the genomic intervals of both anchors and the corresponding local-background test *P* values.

### Contact-distance shift analysis

To quantify the effect of reference choice on inferred genomic contact distances, mappings to each donor-specific haplotype assembly were compared separately with mappings to T2T-CHM13. Shared mapped contact pairs were identified by matching read-pair identifiers between the two mappings. Analyses were restricted to pairs classified as cis in both mappings and assigned to corresponding chromosomes in the two assemblies.

For each retained pair, genomic contact distance was calculated as the absolute difference between the genomic positions of its two mapped read ends. Contact-distance shift was defined as:

ΔD = |D_DSA − D_T2T|,

where D_DSA and D_T2T denote the genomic contact distances of the same read pair in the donor-specific assembly and T2T-CHM13, respectively.

For each haplotype mapping, we summarized the distribution of ΔD and calculated the fraction of retained shared cis contact pairs with distance shifts exceeding 5, 10, 25, 50, or 100 kbp. The denominator for each fraction was the total number of shared cis contact pairs meeting the above criteria in that comparison. These per-haplotype fractions were then summarized across individuals.

### Comparison of loop calls across genome assemblies

Loop calls from each donor-specific haplotype map were compared with those from the T2T-CHM13-based map of the same individual. Within-individual comparisons between haplotypes were performed in both directions, using Hap1 as the query and Hap2 as the target, and vice versa.

Query loop anchors were projected into the target coordinate system using rustybam liftover v0.1.33 (Vollger, 2026) with default parameters and the corresponding assembly-to-assembly alignments in PAF (H. Li, 2018). A projected query anchor was considered matched if it overlapped a target loop anchor interval extended by 10 kbp on each side. All returned projection intervals were considered, and the identifiers of target loops matched by each query anchor were recorded.

Query loops were assigned to three categories: Both anchors, when the two query anchors shared at least one matched target-loop identifier; Single anchor, when at least one query anchor had a match but the two anchors shared no matched target-loop identifier; No anchor, when neither query anchor had a match. Anchors without a returned projection were treated as unmatched, and their query loops were retained in the classification.

### Identification of shared and specific contacts between haplotypes

For each individual, the same Omni-C/Hi-C read pairs were independently mapped to the corresponding Hap1 and Hap2 assemblies. Comparisons used nonduplicate contact pairs generated by the processing workflow described above, with a minimum mapping-quality threshold of 40 applied during contact parsing using pairtools v1.0.3. Read-pair identifiers were compared between the two mapping outputs separately for each sequencing dataset. Pairs whose identifiers were present in both outputs were classified as shared contacts, whereas those present in only one output were initially classified as candidate haplotype-specific contacts.

To minimize specificity arising from differences in duplicate handling between the two mappings, candidate haplotype-specific pairs were further filtered using duplicate records from the alternative haplotype mapping. A candidate was excluded if its read-pair identifier occurred either as a duplicate read ID or as the retained representative of a duplicate group, recorded in the parent_readID field, in the alternative mapping. The remaining candidates were designated haplotype-specific contacts. Their contact-pair records were extracted from the corresponding haplotype mapping outputs and pooled across sequencing datasets within each individual and haplotype for downstream analyses.

### Validation of haplotype-specific contacts

Minimum unique k-mer lengths were determined independently for each assembly using Newmap with --search-range=20:200, and single-read mappability tracks were generated for a read length of 150 bp (Karimzadeh et al., 2018). A contact end was classified as mappable if its genomic position fell within an interval with a single-read mappability score of 1 in the corresponding assembly, indicating coverage by at least one unique 150-mer.

To assess recovery in the alternative haplotype assembly, Hap1-specific read pairs retained after the duplicate-associated filtering described above were extracted from the original FASTQ files using seqtk subseq v1.4 and remapped to the corresponding Hap2 assembly using BWA-MEM v0.7.17 with −5SP -T0. Alignments were processed using pairtools v1.0.3 with --min-mapq 40 -- walks-policy 5unique --max-inter-align-gap 30, followed by sorting and duplicate removal.

### Identification of CDHs

CDHs were identified independently for each individual and each phased haplotype. Haplotype-specific and total contact counts were summarized in non-overlapping 25 kbp windows. Short-range cis contacts and all trans contacts were excluded. Windows with higher contact coverage tend to contain more haplotype-specific contacts, making raw counts unsuitable for direct comparison. We therefore used negative-binomial regression to estimate the expected number of haplotype-specific contacts from the number of shared contacts in each window. The relationship between local contact coverage and haplotype-specific contact counts was allowed to be nonlinear, and chromosome was included as a categorical variable to account for differences in baseline contact counts among chromosomes. For computational efficiency, the model was fitted to a chromosome-stratified random sample of up to 40,000 eligible windows and then applied to all eligible windows. To reduce the influence of extreme high-count windows on background estimation, the model was fitted once, the 0.5% of windows with the largest standardized excesses over the prediction were removed, and the model was fitted a second time. Even after accounting for the coverage-dependent expected count, the remaining variation around the fitted mean may differ across coverage levels. To account for this, windows were divided into 20 quantile-based groups according to their expected counts, and negative-binomial dispersion was estimated separately within each group. Windows in the upper 0.5% of dispersion contributions were excluded when estimating the group-specific dispersion. The resulting estimates were linearly interpolated across groups to obtain a coverage-specific dispersion value for each window. Using the fitted expected count and dispersion, we calculated a one-sided upper-tail *P* value for observing at least the measured number of haplotype-specific contacts. Within each individual-haplotype map, *P* values were adjusted across all tested windows using the Benjamini–Hochberg procedure. A window was classified as a CDH if it had an adjusted q value below 0.01, contained at least twice as many haplotype-specific contacts as expected, and contained at least ten haplotype-specific contacts. Directly adjacent significant windows were merged into a single CDH region.

### Repeat and segmental duplication annotation

Repetitive elements were annotated in each genome assembly using three complementary approaches. Tandem repeats were identified with TRF v4.1.0 using the parameters ‘2 7 7 80 10 50 2000 −l 30 -h -ngs’ (Benson, 1999). Interspersed repeats were annotated with RepeatMasker v4.1.5 using the NCBI engine and the human repeat library with sensitive settings (‘-s -e ncbi -xsmall -species human’). Low-complexity regions were additionally annotated using WindowMasker v2.2.22 in a two-step procedure consisting of count generation followed by interval masking (Morgulis et al., 2006). Repeat annotations from all three approaches were merged, and genome assemblies were soft-masked for downstream analyses.

Segmental duplications were detected on repeat-masked assemblies using SEDEF v1.1 (Numanagić et al., 2018). Candidate duplication pairs greater than 1 kbp in length, sharing greater than 90% sequence identity, and containing less than 70% satellite sequence were retained as segmental duplications. Candidate duplications overlapping Flagger or nucFlag regions were removed to exclude low-confidence assembly regions (Asri et al., 2026). FastCN-based copy number support computed from the corresponding *de novo* assemblies was used as orthogonal validation where applicable.

### SV callsets and filtering

Human SVs were obtained from the HPRC Release 2 SV callset (Lin et al., 2026), and analyses were restricted to insertions and deletions at least 50 bp. For within-individual comparisons, heterozygous SVs were defined as variants present on one haplotype but absent from the other. Each haplotype was assigned carrier or non-carrier status at each locus, and these assignments were used for CDH–SV and gene-expression association analyses. Complex SVs and variants with ambiguous coordinates or insufficient genotype support were excluded from locus-level comparisons.

Nonhuman ape SVs and orthologous sequence states were obtained from the ape T2T genome project (Yoo et al., 2025) and used to infer the ancestral states of human CSA-SVs, as described below.

### Projection of CDHs to T2T-CHM13 and pan-haplotype clustering

CDH intervals identified in DSAs were projected to T2T-CHM13 using precomputed assembly-to-assembly alignments in PAF and rustybam (v0.1.33). For each CDH, projected fragments on the corresponding chromosome were combined into a single interval spanning their minimum start and maximum end coordinates. Cross-chromosome projections were stored separately.

Analyses were restricted to individuals with projected CDHs available for both haplotypes. Overlapping or directly adjacent projected intervals were first merged separately by haplotype and chromosome. CDH recurrence was defined at each genomic position as the number of distinct haplotypes with a projected CDH covering that position, with each haplotype counted once. Contiguous positions meeting the haplotype-support threshold were merged into regions. Short, fragmented regions and regions exceeding 1 Mbp were excluded. Regions supported by at least 20 haplotypes were classified as recurrent CDH regions.

### Identification of CSA-SVs

Associations between SVs and local CDH occurrence were evaluated using haplotype-specific CDH intervals and SV coordinates represented in the T2T-CHM13 coordinate system. Each distinct SV locus was tested separately. For each SV–haplotype combination, local CDH status was defined by whether at least one CDH from that haplotype overlapped a query interval extending 25 kbp (one window of CDH detection) upstream and downstream of the SV position recorded in the input carrier table. Only individuals with both haplotypes represented in the SV and CDH inputs were included.

For insertions, carrier labels were retained on the originally assigned haplotypes. For deletions, carrier labels were exchanged between the two haplotypes of each individual before association testing. These labels defined effective carrier status. SV loci with at least one effective carrier haplotype in the analyzed cohort were eligible for testing. For each locus, a 2 × 2 contingency table was constructed from effective carrier status and local CDH presence. Associations were evaluated using two-sided Fisher’s exact tests on the unmodified counts. Odds ratios were calculated after adding a pseudocount of 1 to each cell: OR = [(a + 1)(d + 1)]/[(b + 1)(c + 1)], where a and b denote effective carrier haplotypes with and without a local CDH, respectively, and c and d denote the corresponding counts among the remaining haplotypes. *P* values were adjusted using the Benjamini–Hochberg procedure across all eligible SV loci, including loci without local CDHs, which were assigned *P* = 1. SVs with OR > 2, an adjusted *P* value < 0.05, and at least three effective carrier haplotypes with a local CDH were designated CSA-SVs.

### Robustness analyses of CSA-SV associations

To evaluate potential confounding by ancestry, local ancestry scores were averaged across the ±25 kbp window surrounding each CSA-SV and included as covariates in a logistic regression model together with standardized genome-wide CDH burden. Local ancestry annotations were obtained from the HPRC Release 2 PCLAI v1.1 tracks in T2T-CHM13 coordinates (Geleta et al., 2026). CDH burden was defined as the total number of CDH intervals per haplotype and standardized to a mean of zero and a standard deviation of one. The association between SV status and local CDH occurrence was compared before and after covariate adjustment.

To assess whether correlation between the two haplotypes from the same individual affected statistical inference, CSA-SV associations were additionally fitted using logistic regression with donor-clustered standard errors, treating the two haplotypes from each individual as correlated observations. Benjamini–Hochberg correction was applied across the tested CSA-SVs.

### Long-read RNA-seq processing

PacBio Kinnex RNA-seq data were analyzed for 163 individuals with matched chromatin contact data. Circular consensus reads were generated using CCS v8.0.1, and concatenated cDNA reads were segmented using Skera v1.2.0. Primer removal and demultiplexing were performed using lima v2.10.0 with --isoseq --peek-guess. Reads were further processed using Iso-Seq refine v4.1.2 with --require-polya to retain poly(A)-containing reads, trim poly(A) tails, and remove residual concatemers, yielding FLNC reads.

FLNC reads were aligned to T2T-CHM13 v2.0 using pbmm2 v1.17.0 with --preset ISOSEQ -- sort -j 12. The resulting coordinate-sorted BAM files were analyzed separately for each individual using IsoQuant v3.10.0 (Prjibelski et al., 2023), with --data_type pacbio_ccs, the T2T-CHM13 v2.0 reference genome, and the CHM13 RefSeq–Liftoff v5.2 annotation. Other IsoQuant parameters were left at their defaults.

### Expression association analysis

Candidate cis genes were selected within 500 kbp of each CSA-SV. Gene expression was quantified using gene-level TPM values from IsoQuant. For each SV–gene pair, individuals were classified as non-carriers (0/0), heterozygous carriers (0/1), or homozygous carriers (1/1). Observations with missing or unrecognized genotypes, missing TPM values, or missing gene identifiers were excluded, as were genes with identifiers beginning with “novel_gene”. Duplicate records were removed so that each individual contributed one observation per SV–gene pair.

SV–gene pairs were tested when at least two genotype classes were represented. Differences in TPM across genotype classes were assessed using Kruskal–Wallis tests. *P* values were adjusted using the Benjamini–Hochberg procedure across all tested SV–gene pairs, and associations with adjusted *P* values below 0.05 were retained as significant expression-associated CSA-SV–gene pairs.

### Aggregated insulation analysis

For selected CSA-SV locus, individuals with nonmissing genotypes were grouped as 0/0, 0/1, or 1/1. Local contact matrices were extracted from contact maps at 25 kbp resolution in T2T-CHM13 coordinates using cooler (version 0.9.3). Extraction windows extended from 500 kbp upstream of the SV start to 500 kbp downstream of the analysis interval end, defined as the SV start plus the reported SV length. Each individual matrix was divided by the sum of its entries, and the resulting matrices were averaged within each genotype group. Groups containing at least two individuals with valid matrices were retained. Genotype differences were visualized as pixelwise log ratios.

Insulation profiles were generated separately for each locus using precomputed per-individual score tracks from the TAD analysis described above. Scores were extracted at 25 kbp intervals across the same genomic windows, with positions expressed relative to the SV start. At each position, the mean and standard error of the mean were calculated across individuals within each genotype group, excluding missing scores.

### Insulation-expression correlation analysis

To assess the relationship between local chromatin insulation and transcription, insulation scores at CSA-SV-associated loci were correlated with expression levels of the corresponding cis genes across matched samples. Only samples with valid measurements for both insulation and gene expression were included. Associations were evaluated using two-sided Spearman rank correlation, with multiple-testing correction using the Benjamini–Hochberg procedure where applicable.

### Inference of ancestral and human-lineage-derived CSA-SVs

Ancestral states of CSA-SVs were inferred using T2T assemblies from six nonhuman ape species, including chimpanzee, bonobo, gorilla, Bornean orangutan, Sumatran orangutan, and siamang. For each human CSA-SV, the corresponding orthologous interval was identified in nonhuman ape assemblies using whole-genome alignments together with flanking sequence support. Only loci with confident orthology, unique placement, and sufficient alignment coverage were retained for ancestral-state inference. The allelic state of each CSA-SV was determined by comparing the human SV allele with the sequence state of the orthologous ape interval. CSA-SVs were classified as ancestral, or ape-matching, when the SV allele matched the orthologous allelic state in at least one nonhuman ape haplotype. CSA-SVs were classified as human-lineage derived when the SV allele was absent from all confidently aligned nonhuman ape orthologous intervals. Loci with ambiguous alignments, conflicting allele representations, multiple equally supported placements, or insufficient sequence support were excluded from ancestral-state assignment.

### Human–ape chromatin contact divergence analysis

Local chromatin contact patterns were compared between humans and chimpanzees at a representative 2.7 kbp human-derived deletion at chr1:88,853,624–88,856,340 (T2T-CHM13). Contact maps were generated separately for human individual HG01786, homozygous for the derived deletion allele, and a chimpanzee carrying the inferred ancestral, undeleted allele. Local contact matrices were examined at 10 kbp resolution within orthologous regions corresponding to 500 kbp upstream and downstream of the deletion start. Both maps were displayed in T2T-CHM13 coordinates to compare local chromatin contact patterns surrounding the deletion.

### PheWAS

PheWAS was performed using Firth logistic regression implemented in PLINK 2.0 (Chang et al., 2015), which reduces small-sample bias and provides finite estimates in the presence of separation (Wang, 2014; Ma et al., 2013). SV genotypes were obtained using Locityper (Prodanov et al., 2025). For comparison with nearby sequence variants, single-nucleotide polymorphisms (SNPs) within 100 kbp of each SV breakpoint were extracted from the *All of Us* short-read variant call set (Bick et al., 2024), normalized to biallelic variants using BCFtools (Danecek et al., 2021), and tested for linkage disequilibrium with the corresponding SV using PLINK 2.0.

Phecodes were derived from ICD-10-coded electronic health records (Wu et al., 2019). Cases were required to have at least two occurrences of the corresponding phecode, whereas participants with no occurrence were classified as controls. Phenotypes with fewer than 20 cases were excluded, resulting in 1,621 phenotypes for analysis. Related individuals were excluded, and association models included age, sex, and the first ten ancestry principal components as covariates. Analyses were performed separately for each SV and flanking SNP using individuals with valid genotypes for the tested variant.

To limit associations supported by sparse genotype counts, we retained results only when at least five cases carrying the tested allele and genotype data were available for at least 200,000 participants. *P* values were corrected for testing across phenotypes using the multiple-testing correction. PheWAS plots were generated using pheTK (v0.2.12) (Tran et al., 2025).

### Statistics and reproducibility

Statistical analyses were performed in R unless otherwise specified. Comparisons between two independent groups used two-sided Wilcoxon rank-sum tests, whereas paired comparisons used two-sided Wilcoxon signed-rank tests. Differences in gene expression across SV genotype groups were assessed using Kruskal–Wallis tests. Associations between SV status and CDH presence were evaluated using two-sided Fisher’s exact tests on 2 × 2 contingency tables, with odds ratios reported as measures of association. CDH detection used one-sided upper-tail tests under the negative-binomial model. Permutation-based empirical tests were used for recurrence and enrichment analyses, following the randomization procedures described in the corresponding Methods subsections.

Multiple-testing correction was performed using the Benjamini–Hochberg procedure, with the scope of correction specified for each analysis. For the selected *GBP3* locus, the expression comparison used the adjusted *P* value from the original expression-association scan. For box plots, center lines indicate medians, boxes span the IQR, and whiskers extend to the most extreme observations within 1.5 times the IQR below the first quartile or above the third quartile.

## ACKNOWLEDGMENTS

We thank Tonia Brown for editing the manuscript and supplementary materials. Research reported in this publication was supported, in part, by the U.S. National Human Genome Research Institute of the National Institutes of Health (NIH) (U01HG010971, 1U41HG007497, R01HG002385, R01HG010169 to E.E.E.). The content is solely the responsibility of the authors and does not necessarily represent the official views of the NIH. E.E.E. is an investigator of the Howard Hughes Medical Institute.

## AUTHOR CONTRIBUTIONS

E.E.E. and L.N. conceptualized the study. L.N., J.L., and D.Y. conducted formal analyses and created visualizations. L.N. and E.E.E. did the interpretation. L.N. and E.E.E. wrote the original draft. E.E.E., W.S.N., J.M., and X.S. supervised the study. All authors reviewed and edited the manuscript.

## DECLARATION OF GENERATIVE AI AND AI-ASSISTED TECHNOLOGIES IN THE MANUSCRIPT PREPARATION PROCESS

During the preparation of this work, ChatGPT and Claude were used for code improvement and text refinement. After using these tools, the authors reviewed and edited the content as needed and take full responsibility for the content of the published article.

## RESOURCE AVAILABILITY MATERIALS AVAILABILITY

This study did not generate new unique reagents.

## DATA AVAILABILITY

Human genome assembly data, Hi-C/Omni-C data, and transcriptome data used in this study are available through AnVIL and the AWS Open Data Program. Detailed data-access information, including sample metadata, accession numbers, and download paths, is provided in the HPRC Release 2 data tables: https://github.com/human-pangenomics/hprc_intermediate_assembly/tree/main/data_tables

Nonhuman ape data used in this study are available through NCBI and GenomeArk. Accession numbers and download links are provided in the T2T-Primates project repository: https://github.com/marbl/Primates

## CODE AVAILABILITY

Custom scripts used for the analyses and figure generation are available at GitHub (https://github.com/LingbinNi/pangenome_chromatin_associated_SVs). Software versions and analysis parameters are described in Methods.

## LEAD CONTACT

Requests for further information and resources should be directed to and will be fulfilled by the lead contact, Evan E. Eichler.

**Human Pangenome Reference Consortium Version 2 Authors**

Derek Albracht^1^, Ivan A. Alexandrov^2^, Jamie Allen^3^, Alawi A. Alsheikh-Ali^4^, Nicolas Altemose^5^, Casey Andrews^6^, Dmitry Antipov^7^, Lucinda Antonacci-Fulton^1^, Alexander Arguello^8^, Mobin Asri^9^, Marcelo Ayllon^10^, Jennifer R. Balacco^11^, Floris P. Barthel^12^, Edward A. Belter Jr^1^, Halle D. Bender^9^, Andrew P. Blair^9^, Davide Bolognini^13^, Katherine E. Bonini^14^, Christina Boucher^15^, Guillaume Bourque^16,17,18^, Silvia Buonaiuto^19^, Shuo Cao^19^, Andrew Carroll^20^, Ann M. Mc Cartney^21^, Monika Cechova^9^, Mark J.P. Chaisson^22^, Pi-Chuan Chang^20^, Xian Chang^9^, Jitender Cheema^3^, Haoyu Cheng^23^, Claudio Ciofi^24^, Hiram Clawson^9^, Sarah Cody^1^, Vincenza Colonna^19^, Holland C. Conwell^25^, Robert Cook-Deegan^26^, Mark Diekhans^9^, Maria Angela Diroma^24^, Daniel Doerr^27,28,29^, Zheng Dong^6^, Danilo Dubocanin^5^, Richard Durbin^30,31^, Jana Ebler^27,32^, Evan E. Eichler^10,33^, Jordan M. Eizenga^9^, Parsa Eskandar^9^, Eddie Ferro^15^, Anna-Sophie Fiston-Lavier^34,35^, Sarah M. Ford^25^, Willard W. Ford^36^, Giulio Formenti^11^, Adam Frankish^3^, Mallory A. Freeberg^3^, Qichen Fu^6^, Stephanie M. Fullerton^37^, Robert S. Fulton^1^, Shenghan Gao^38^, Yan Gao^39^, Gage H. Garcia^10^, Obed A. Garcia^40^, Joshua M.V. Gardner^9^, Shilpa Garg^41^, Erik Garrison^19^, Nanibaa’ A. Garrison^42,43,44^, John E. Garza^1^, Margarita Geleta^45,46^, Mohammadmersad Ghorbani^47^, Tina A. Graves-Lindsay^1^, Richard E. Green^25^, Carol W. Greider^48^, Cristian Groza^49^, Bida Gu^22^, Andrea Guarracino^12,19^, Melissa Gymrek^50^, Maximilian Haeussler^9^, Leanne Haggerty^3^, Ira M. Hall^51,52^, Nancy F. Hansen^7^, Yue Hao^12^, Mohammad Amiruddin Hashmi^4^, David Haussler^9^, Prajna Hebbar^9^, Peter Heringer^27,28,29^, Glenn Hickey^9^, Todd L. Hillaker^9^, S. Nakib Hossain^3^, Neng Huang^39,53^, Sarah E. Hunt^3^, Toby Hunt^3^, Alexander G. Ioannidis^5,9,46^, Nafiseh Jafarzadeh^9^, Nivesh Jain^11^, Erich D. Jarvis^11,33^, Maryam Jehangir^12^, Juan Jiang^6^, Eimear E. Kenny^14^, Juhyun Kim^7^, Bonhwang Koo^11^, Sergey Koren^7^, Milinn Kremitzki^1,6^, Charles H. Langley^54^, Ben Langmead^55^, Heather A. Lawson^6^, Daofeng Li^6^, Heng Li^39,53^, Ronghan Li^6^, Wen-Wei Liao^51,52^, Jiadong Lin^10^, Tianjie Liu^6^, Glennis A. Logsdon^38^, Ryan Lorig-Roach^9^, Jonathan LoTempio Jr^21,56^, Hailey Loucks^9^, Jane E. Loveland^3^, Jianguo Lu^57^, Shuangjia Lu^51,52^, Julian K. Lucas^9^, Walfred Ma^22^, Juan F. Macias-Velasco^1,6,58^, Kateryna D. Makova^59^, Maximillian G. Marin^39,53^, Christopher Markovic^1^, Tobias Marschall^27,32^, Franco L. Marsico^19^, Fergal J. Martin^3^, Mira Mastoras^9^, Capucine Mayoud^34^, Brandy McNulty^9^, Jack A. Medico^11^, Julian M. Menendez^9^, Karen H. Miga^9^, Anna Minkina^60^, Matthew W. Mitchell^61^, Saswat K. Mohanty^62^, Younes Mokrab^47,63,64^, Jean Monlong^65^, Shabir Moosa^47^, Avelina Moreno-Ochando^66,67^, Shinichi Morishita^68^, Jonathan M. Mudge^3^, Katherine M. Munson^10^, Njagi Mwaniki^69^, Nasna Nassir^4^, Chiara Natali^24^, Shloka Negi^9^, Lingbin Ni^10^, Adam M. Novak^9^, Faith Okamoto^9^, Keisuke K. Oshima^38^, Pilar N. Ossorio^70,71^, Chie Owa^68^, Sadye Paez^11^, Benedict Paten^9^, Clelia Peano^13,72^, Adam M. Phillippy^7,55,73,74^, Brandon D. Pickett^7^, Laura Pignata^19^, Nadia Pisanti^69^, David Porubsky^10,75^, Pjotr Prins^19^, Timofey Prodanov^27,32^, Anandi Radhakrishnan^9^, T. Rhyker Ranallo-Benavidez^12^, Brian J. Raney^9^, Mikko Rautiainen^76^, Alessandro Raveane^13^, Andreas Rechtsteiner^48^, Luyao Ren^10,33^, Arang Rhie^7^, Fedor Ryabov^77,78^, Samuel Sacco^25^, Farnaz Salehi^19^, Michael C. Schatz^55,79^, Laura B. Scheinfeldt^80^, Aarushi Sehgal^36^, William E. Seligmann^25^, Mahsa Shabani^81^, Kishwar Shafin^20^, Shadi Shahatit^34^, Ruhollah Shemirani^14^, Vikram S. Shivakumar^55^, Swati Sinha^3^, Jouni Sirén^9^, Linnéa Smeds^62^, Steven J. Solar^7^, Marco Sollitto^11,24^, Nicole Soranzo^13,30,82^, Andrew B. Stergachis^10,60^, Marie-Marthe Suner^3^, Yoshihiko Suzuki^68^, Arda Söylev^27,32^, Ahmad Abou Tayoun^83,84^, Jack A.S. Tierney^3^, Chad Tomlinson^1^, Francesca Floriana Tricomi^3^, Mohammed Uddin^4,85^, Matteo Tommaso Ungaro^25,86^, Rahul Varki^15^, Flavia Villani^19^, Ivo Violich^9^, Mitchell R. Vollger^87^, Brian P. Walenz^7^, Charles Wang^88^, Lisa E. Wang^14^, Ting Wang^1,6,58^, Aaron M. Wenger^89^, Conor V. Whelan^11^, Zilan Xin^6^, Zheng Xu^6^, Kai Ye^90^, DongAhn Yoo^10^, Wenjin Zhang^6^, Ying Zhou^39^, Xiaoyu Zhuo^6^, Giulia Zunino^13^

**Affiliations**

^1^McDonnell Genome Institute, Washington University School of Medicine, St. Louis, MO 63108, USA

^2^Department of Human Molecular Genetics and Biochemistry, Faculty of Medical and Health Sciences, Tel Aviv University, Tel Aviv 69978, Israel

^3^European Molecular Biology Laboratory, European Bioinformatics Institute (EMBL-EBI), Wellcome Genome Campus, Hinxton, Cambridge CB10 1SD, UK

^4^Center for Applied and Translational Genomics (CATG), Mohammed Bin Rashid University of Medicine and Health Sciences, Dubai Health, Dubai, UAE

^5^Department of Genetics, Stanford University, Palo Alto, CA 94304 USA

^6^Department of Genetics, Washington University School of Medicine, St. Louis, MO 63110, USA

^7^Genome Informatics Section, Center for Genomics and Data Science Research, National Human Genome Research Institute, National Institutes of Health, Bethesda, MD 20892, USA

^8^Division of Genome Sciences, National Human Genome Research Institute, Bethesda, MD 20871 USA

^9^UC Santa Cruz Genomics Institute, University of California, Santa Cruz, CA 95060, USA

^10^Department of Genome Sciences, University of Washington School of Medicine, Seattle, WA 98195, USA

^11^The Vertebrate Genome Laboratory, The Rockefeller University, New York, NY 10065, USA

^12^Bioinnovation and Genome Sciences, The Translational Genomics Research Institute (TGen), Phoenix, AZ 85004, USA

^13^Human Technopole, Milan, Italy

^14^Institute for Genomic Health, Icahn School of Medicine at Mount Sinai, New York, NY 10029, USA

^15^Department of Computer and Information Science and Engineering, University of Florida, Gainesville, FL 32611, USA

^16^Canadian Center for Computational Genomics, McGill University, Montréal, QC H3A 0G1, Canada

^17^Department of Human Genetics, McGill University, Montréal, QC H3A 0G1, Canada

^18^Victor Phillip Dahdaleh Institute of Genomic Medicine, Montréal, QC H3A 0G1, Canada

^19^Department of Genetics, Genomics and Informatics, University of Tennessee Health Science Center, Memphis, TN 38163, USA

^20^Google LLC, Mountain View, CA 94043, USA

^21^Institute of Clinical and Translational Sciences, University of California, Irvine, CA 92697, USA

^22^Quantitative and Computational Biology, University of Southern California, Los Angeles, CA 90089, USA

^23^Department of Biomedical Informatics and Data Science, Yale School of Medicine, New Haven, CT 06510, USA

^24^Department of Biology, University of Florence, Sesto Fiorentino, FI 50019, Italy

^25^Department of Ecology and Evolutionary Biology, University of California, Santa Cruz, CA 95060, USA

^26^Arizona State University, Consortium for Science, Policy & Outcomes, Washington, DC 20006, USA

^27^Center for Digital Medicine, Heinrich Heine University Düsseldorf, Düsseldorf, NRW, DE

^28^Department for Endocrinology and Diabetology at the Medical Faculty and University Hospital Düsseldorf, Heinrich Heine University Düsseldorf, Düsseldorf, NRW, DE

^29^Paul-Langerhans-Group Computational Diabetology, German Diabetes Center (DDZ) and Leibniz Institute for Diabetes Research, Düsseldorf, NRW, DE

^30^Wellcome Sanger Institute, Genome Campus, Hinxton, CB10 1RQ, UK

^31^Department of Genetics, University of Cambridge, Cambridge, CB2 3EH, UK

^32^Institute for Medical Biometry and Bioinformatics, Medical Faculty and University Hospital Du sseldorf, Heinrich Heine University, Du sseldorf, NRW, DE

^33^Howard Hughes Medical Institute, Chevy Chase, MD 20815, USA

^34^ISEM, Univ Montpellier, CNRS, IRD, Montpellier, FR

^35^Institut Universitaire de France, Paris, FR

^36^Department of Computer Science and Engineering, University of California San Diego, La Jolla, CA 92093, USA

^37^Department of Bioethics & Humanities, University of Washington School of Medicine, Seattle, WA 98195, USA

^38^Department of Genetics, Epigenetics Institute, Perelman School of Medicine, University of Pennsylvania, Philadelphia, PA 19104, USA

^39^Department of Data Science, Dana-Farber Cancer Institute, Boston, MA 02215, USA

^40^Department of Anthropology, University of Kansas, Lawrence, KS 66045, USA

^41^School of Health Sciences, University of Manchester, Manchester M13 9PL, UK

^42^Traditional, ancestral and unceded territory of the Gabrielino/Tongva peoples, Institute for Society & Genetics, University of California, Los Angeles, Los Angeles, CA 90095, USA

^43^Traditional, ancestral and unceded territory of the Gabrielino/Tongva peoples, Institute for Precision Health, David Geffen School of Medicine, University of California, Los Angeles, Los Angeles, CA 90095, USA

^44^Traditional, ancestral and unceded territory of the Gabrielino/Tongva peoples, Division of General Internal Medicine & Health Services Research, David Geffen School of Medicine, University of California, Los Angeles, Los Angeles, CA 90095, USA

^45^Department of Electrical Engineering and Computer Science, University of California, Berkeley, Berkeley, CA 94720, USA

^46^Department of Biomedical Data Science, Stanford University School of Medicine, Stanford, CA 94305, USA

^47^Medical and Population Genomics Lab, Sidra Medicine, Doha, Qatar

^48^Department of Molecular Cell and Developmental Biology, University of California, Santa Cruz, CA, USA

^49^Montreal Heart Institute, Montréal, QC, Canada

^50^Department of Pediatrics, University of California San Diego, La Jolla, CA 92093, USA

^51^Center for Genomic Health, Yale University School of Medicine, New Haven, CT 06510, USA

^52^Department of Genetics, Yale University School of Medicine, New Haven, CT 06510, USA

^53^Department of Biomedical Informatics, Harvard Medical School, Boston, MA 02115, USA

^54^Department of Evolution and Ecology and the Center for Population Biology, University of California, One Shields, Davis, CA 95616, USA

^55^Department of Computer Science, Johns Hopkins University, Baltimore, MD 21218, USA

^56^Department of Pediatrics, Division of Genetics, School of Medicine, University of California, Irvine, CA 92697, USA

^57^Sun Yat-sen University, Guangzhou, China

^58^Edison Family Center for Genome Sciences & Systems Biology, Washington University School of Medicine, St. Louis, MO 63110, USA

^59^Department of Biology and Center for Medical Genomics, Penn State University, University Park, PA 16802, USA

^60^Division of Medical Genetics, Department of Medicine, University of Washington School of Medicine, Seattle, WA 98195, USA

^61^The Jackson Laboratory for Genomic Medicine, Farmington, CT 06032, USA

^62^Department of Biology, Penn State University, University Park, PA 16802, USA

^63^Department of Biomedical Science, College of Health Sciences, Qatar University, Doha, Qatar

^64^Department of Genetic Medicine, Weill Cornell Medicine-Qatar, Doha, Qatar

^65^IRSD - Digestive Health Research Institute, University of Toulouse, INSERM, INRAE, ENVT, UPS, Toulouse, FR

^66^MATCH biosystems, S.L., Elche, Spain

^67^Universidad Miguel Hernández de Elche, Elche, Spain

^68^Department of Computational Biology and Medical Sciences, The University of Tokyo, Kashiwa, Chiba 277-8561, Japan

^69^Department of Computer Science, University of Pisa, Pisa, Italy

^70^Law School, University of Wisconsin-Madison, Madison, WI 53706, USA

^71^Morgridge Institute for Research, Madison, WI 53715, USA

^72^Institute of Genetics and Biomedical Research, UoS of Milan, National Research Council, Milan, Italy

^73^Department of Biomedical Engineering, Johns Hopkins University, Baltimore, MD 21218, USA

^74^Department of Genetic Medicine, Johns Hopkins University School of Medicine, Baltimore, MD 21205, USA

^75^Genome Biology Unit, European Molecular Biology Laboratory (EMBL), Heidelberg, DE

^76^Institute for Molecular Medicine Finland, Helsinki Institute of Life Science, University of Helsinki, Helsinki, Finland

^77^The Center for Bio- and Medical Technologies, Moscow, RUS

^78^Centre for Biomedical Research and Technology, HSE University, Moscow, RUS

^79^Department of Biology, Johns Hopkins University, Baltimore, MD 21218, USA

^80^Coriell Institute for Medical Research, Camden, NJ 08103, USA

^81^University of Amsterdam, Amsterdam, Netherlands

^82^School of Clinical Medicine, University of Cambridge, Cambridge, CB2 0SP, UK

^83^Center for Genomic Discovery, Mohammed Bin Rashid University, Dubai Health, UAE

^84^Dubai Health Genomic Medicine Center, Dubai Health, UAE

^85^GenomeArc Inc, Mississauga, ON, Canada

^86^Department of Biology and Biotechnologies “Charles Darwin”, University of Rome “La Sapienza”, Rome 00185, IT

^87^Department of Human Genetics and Utah Center for Genetic Discovery, University of Utah, Salt Lake City, UT, USA

^88^Center for Genomics, Loma Linda University School of Medicine, Loma Linda, CA 92350, USA

^89^PacBio, Menlo Park, CA 94025, USA

^90^The first affiliated hospital of Xi’an Jiaotong University, Xi’an Jiaotong University, Xi’an, Shaanxi, 710049, China

