## Supplementary Information for "Structural variants from donor-specific assemblies affect recurrent chromatin contact divergence in humans"

### SUPPLEMENTAL INFORMATION

### SUPPLEMENTAL FIGURES

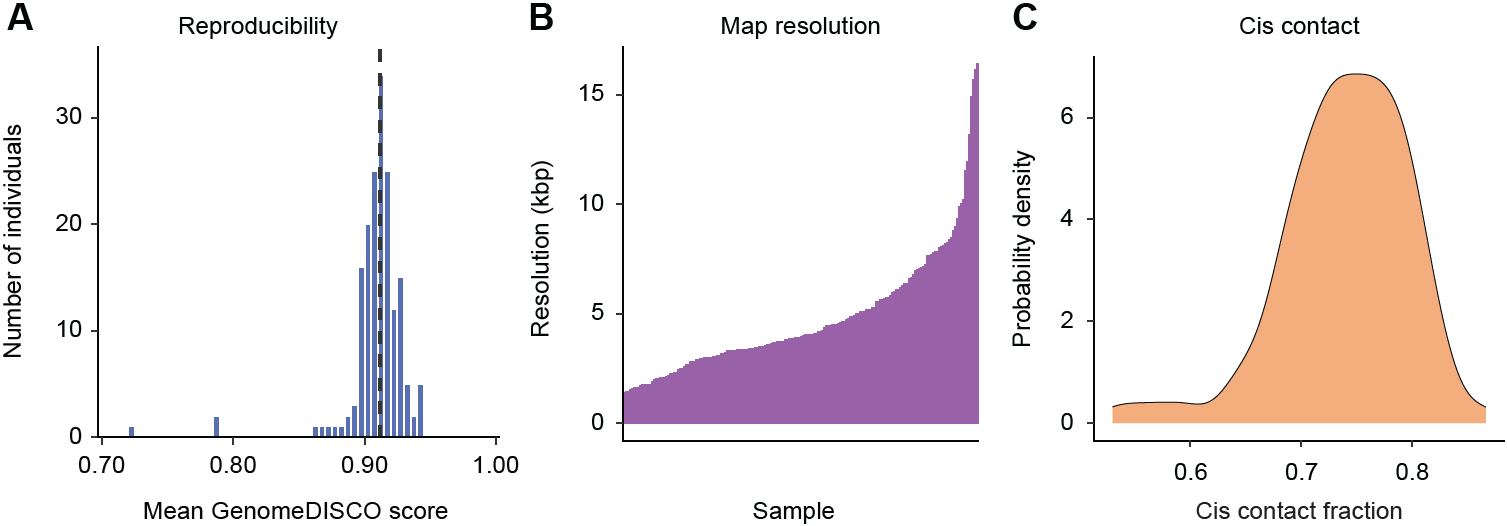

**Figure S1. Quality control of population-scale chromatin contact maps, related to Figure 1**

(A) Distribution of sample-level mean GenomeDISCO reproducibility scores between replicate libraries. Each individual contributes one value. The dashed vertical line indicates the median across individuals.

(B) Estimated contact-map resolution for each sample, ordered by increasing resolution. Resolution was defined as the smallest bin size where at least 80% of genomic bins contained at least 1,000 contacts.

(C) Density distribution of cis-interaction fractions across 177 samples. Cis-interaction fraction was calculated as the proportion of intrachromosomal contact pairs among all contact pairs.

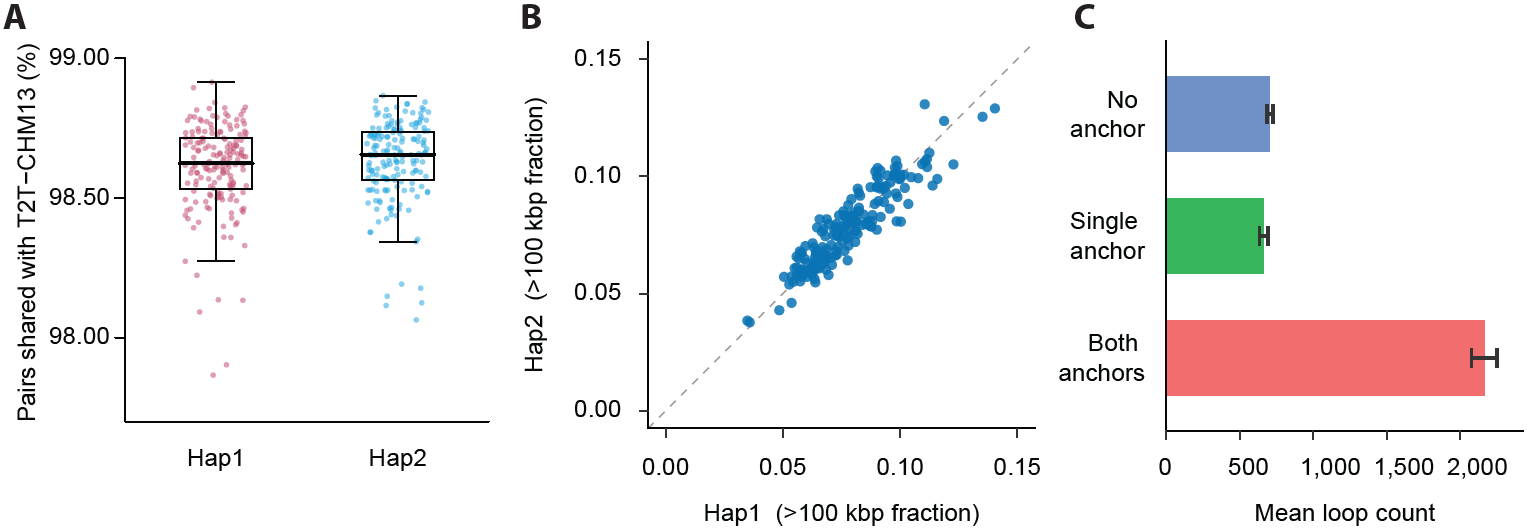

**Figure S2. Concordance of chromatin contacts and loop calls across haplotypes and reference coordinate systems, related to Figure 1**

(A) Percentage of contact pairs recovered by DSA mapping that are also recovered by T2T-CHM13 mapping, shown separately for Hap1 and Hap2. Each point represents one haplotype from one individual. Boxes indicate the IQR, center lines denote medians, and whiskers extend to the most extreme values within 1.5 × IQR of the box.

(B) Comparison between Hap1 and Hap2 of the fractions of shared cis contacts whose inferred genomic distances differ by more than 100 kbp between DSA-based and T2T-CHM13-based mappings. Each blue point represents one individual. The dashed line indicates the identity line.

(C) Mean numbers of DSA-derived loops classified according to their anchor-level concordance with loops identified in the corresponding T2T-CHM13-based maps. Loops were classified as matching at both anchors, at a single anchor, or at neither anchor. Bars indicate the mean across individuals after averaging Hap1 and Hap2 within each individual; error bars denote s.e.m.

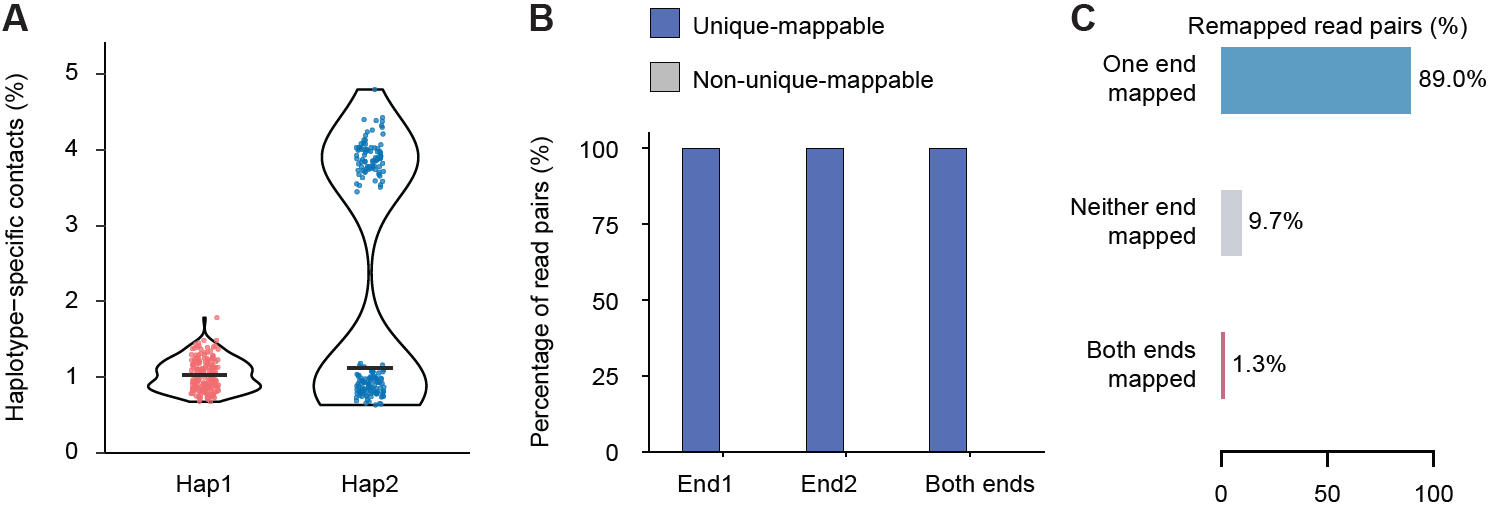

**Figure S3. Validation and characterization of haplotype-specific contacts, related to Figure 2**

(A) Percentages of haplotype-specific contacts in Hap1 and Hap2 across individuals. Each point represents one individual. Percentages were calculated separately for each haplotype as the number of haplotype-specific contacts divided by the sum of shared and haplotype-specific contacts. Violin plots show the distributions, and horizontal bars indicate medians. The two modes in Hap2 correspond to male and female samples and reflect differences in the contribution of X-chromosome–associated contacts.

(B) Mappability assessment of haplotype-specific Hi-C contacts in a representative HG002 Hap1. Bars show the fractions of uniquely mappable and non-uniquely mappable read ends for end 1, end 2, and complete contact pairs with both ends uniquely mappable.

(C) Mapping outcomes of haplotype-specific read pairs remapped to the alternate haplotype assembly. Bars show the percentages of read pairs with only one end mapped (89.0%), neither end mapped (9.7%), or both ends mapped (1.3%). Percentages were calculated from pooled contact counts.

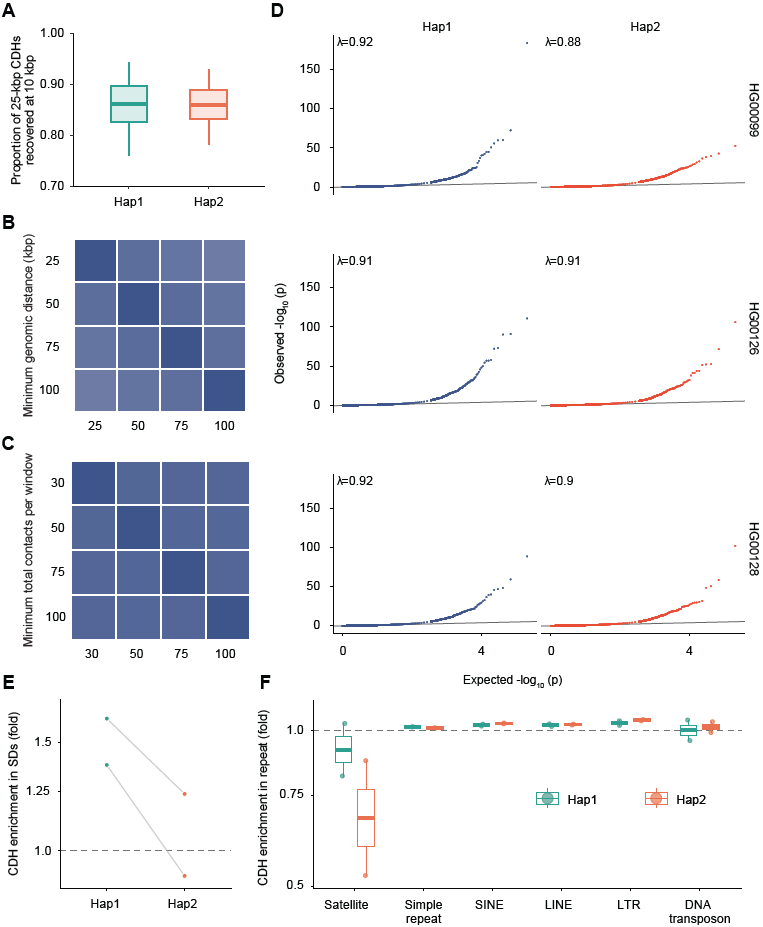

**Figure S4. Robustness, calibration, and genomic context of CDH detection, related to Figure 2**

(A) Recovery of CDHs identified using 25 kbp windows by those identified using 10 kbp windows. Recovery was defined as overlap between merged CDH regions. Box plots show recovery proportions separately for Hap1 and Hap2. Center lines indicate medians, boxes indicate IQRs, and whiskers extend to the most extreme observations within 1.5 × IQR of the box.

(B) Robustness of CDH identification to the minimum genomic distance used to define long-range chromatin contacts. Heatmap colors indicate pairwise Jaccard indices between CDH sets identified using minimum distance thresholds of 25, 50, 75 and 100 kbp.

(C) Robustness of CDH identification to minimum contact-coverage thresholds. Heatmap colors indicate pairwise Jaccard indices between CDH sets identified using minimum total-contact thresholds of 30, 50, 75 and 100 contacts per window.

(D) Quantile–quantile plots of the raw, one-sided negative-binomial *P* values used for CDH detection. Genomic inflation factors (λ) are shown in each panel. Gray lines indicate agreement between observed and expected *P* values. The close agreement between observed and expected values across most of the distribution indicates no systematic inflation of the test statistic, whereas deviations in the upper tail reflect windows with excess haplotype-specific contacts.

(E) Enrichment of CDHs within SDs. Points represent individual haplotypes, and gray lines connect Hap1 and Hap2 from the same individual. The dashed horizontal line indicates the expectation from matched randomized regions.

(F) Enrichment of CDHs across major repeat classes. Box plots show the median and IQR across haplotypes, with individual haplotypes overlaid. The dashed horizontal line indicates the expectation from matched randomized regions. SINE, short interspersed nuclear element; LINE, long interspersed nuclear element; LTR, long terminal repeat.

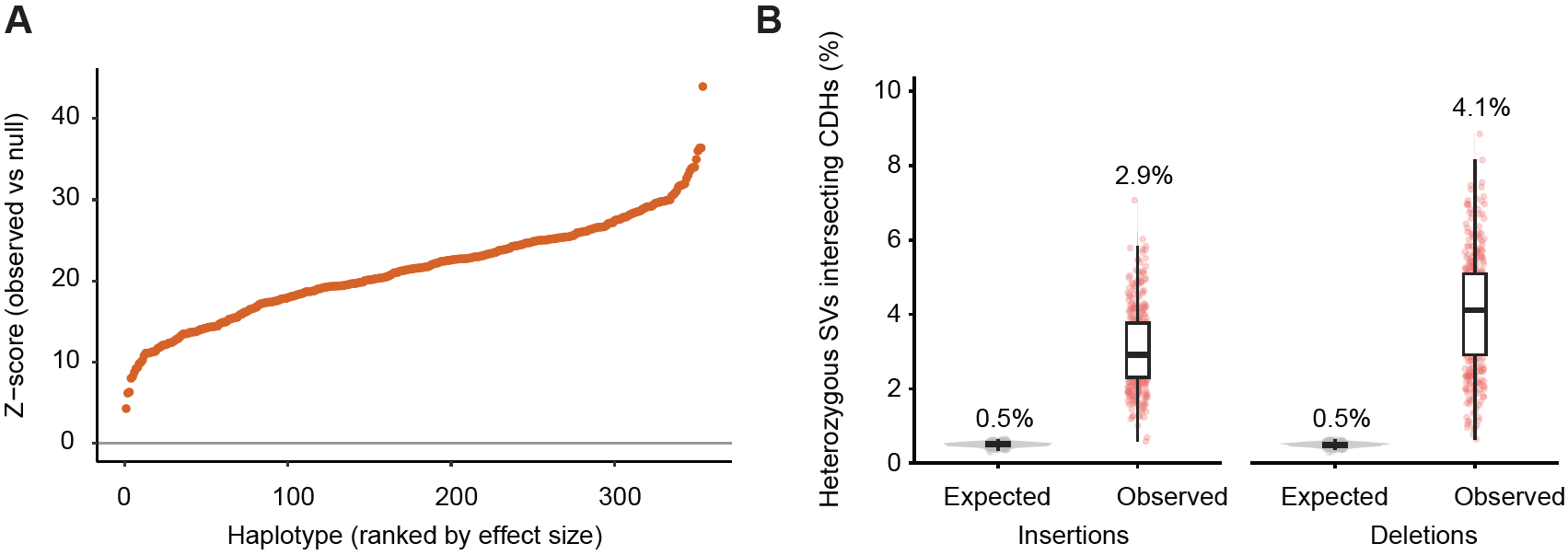

**Figure S5. Enrichment of heterozygous SV overlap with CDHs across haplotypes, related to Figure 2**

(A) Z scores quantifying the overlap of heterozygous SVs with CDHs across haplotypes. For each haplotype, the observed number of SVs intersecting at least one CDH was compared with a null distribution generated from 1,000 chromosome-constrained permutations of CDH intervals that preserved chromosome assignment, interval number, and interval length. Each point represents one haplotype; haplotypes are ordered by increasing Z score.

(B) Observed and expected percentages of heterozygous insertion breakpoints and deletion intervals intersecting CDHs across haplotypes. Expected percentages were calculated from the permutation-derived null distributions described in (A). Points represent individual haplotypes. Boxes show the median and IQR, and whiskers extend to 1.5 times the IQR. Values above the boxes indicate medians.

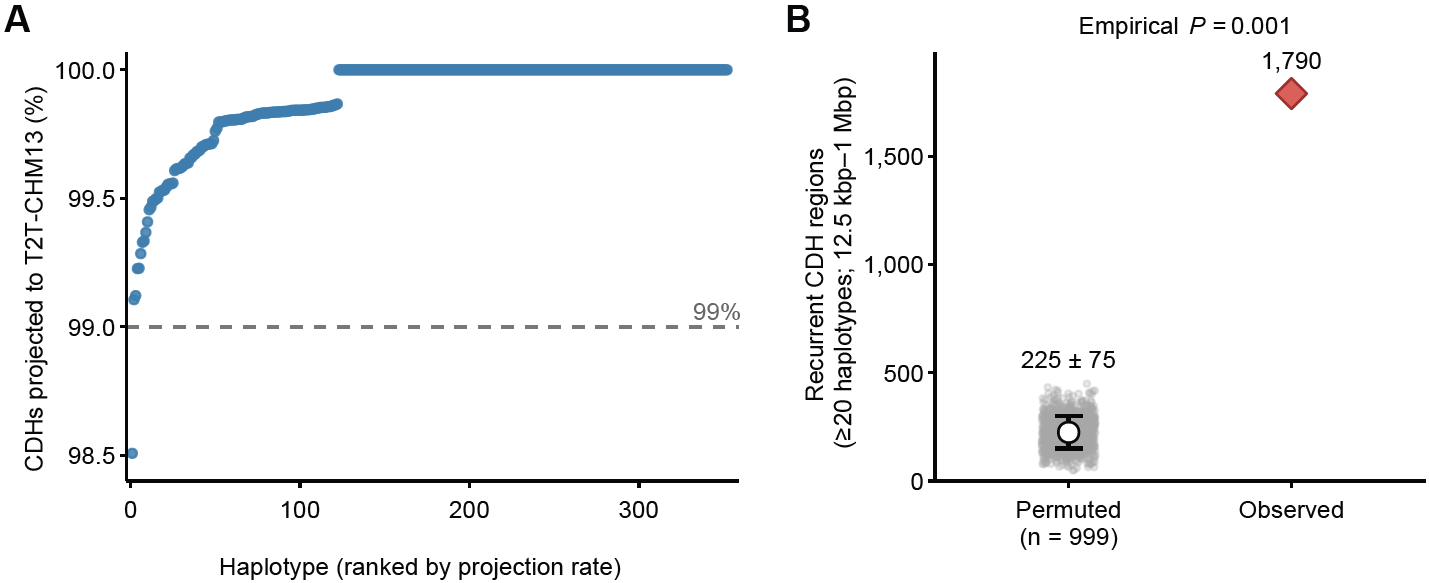

**Figure S6. Projection and recurrence of CDHs across haplotypes, related to Figure 3**

(A) Percentage of CDHs successfully projected from each DSA to T2T-CHM13. Each point represents one haplotype, with haplotypes ordered by increasing projection rate. The dashed line indicates a projection rate of 99%.

(B) Observed number of recurrent CDH regions on autosomes compared with the null distribution obtained from 999 permutations. Gray points represent individual permutations; the open circle and error bar indicate the permutation mean ± s.d.; and the red diamond indicates the observed count. The one-sided empirical *P* value was calculated as (1 + the number of permuted counts greater than or equal to the observed count)/(1 + the number of permutations), yielding *P* = 0.001.

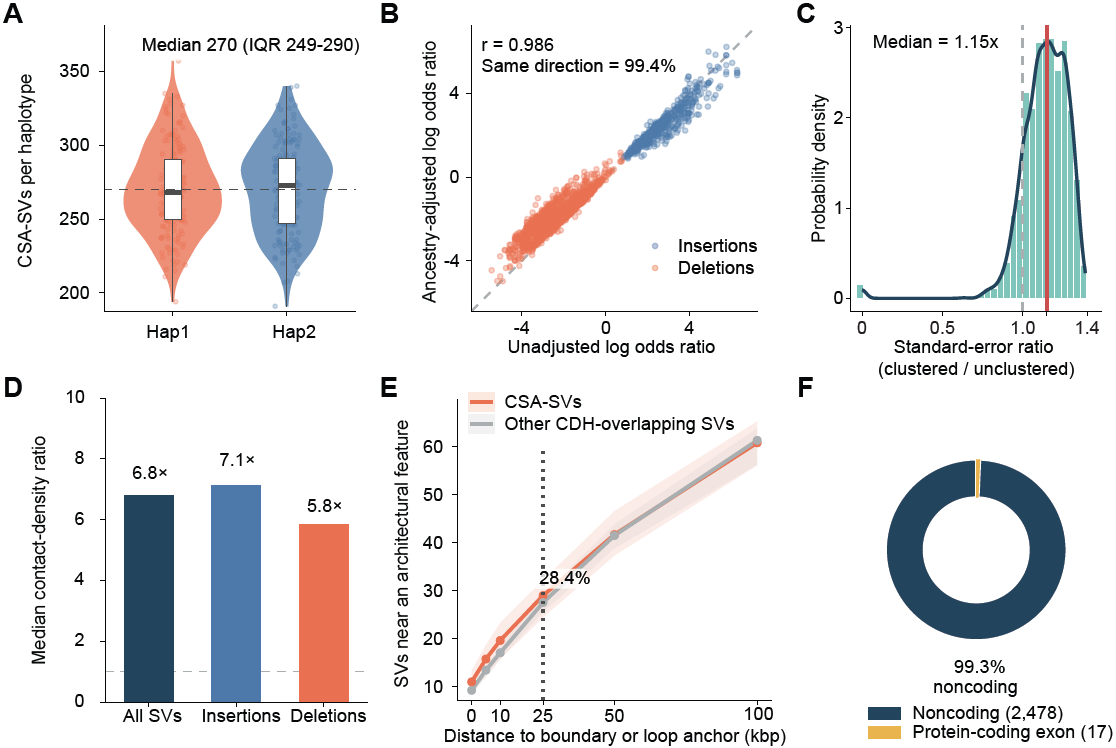

**Figure S7. Robustness and genomic characteristics of CSA-SVs, related to Figure 3**

(A) Distribution of the number of CSA-SVs carried by each haplotype. Data are shown separately for Hap1 and Hap2 across haplotypes. The dashed line indicates the overall median of 270 CSA-SVs per haplotype (IQR, 249–290). Violins show the distributions, boxes indicate medians and IQRs, and whiskers extend to 1.5 times the IQR.

(B) Comparison of CSA-SV effect estimates before and after adjustment for local genetic ancestry. Each point represents one tested SV with a stable model fit; insertions and deletions are indicated by color. The dashed line indicates the identity line. Effect estimates were highly correlated before and after adjustment (Pearson’s r = 0.986), and 99.4% retained the same direction of effect.

(C) Changes in the standard errors of SV effect estimates after accounting for correlation between the two haplotypes from each individual. The histogram shows the ratio of standard errors calculated with versus without clustering by individual. The dashed line indicates no change (ratio = 1), and the solid line indicates the median ratio of 1.15.

(D) Median ratios of local contact density around CSA-SVs relative to other CDH-overlapping SVs. Within each individual, the ratio was calculated by dividing the median contact density around CSA-SVs by that around the comparator SVs. Bars show medians with the ratios: 6.8 for all SVs, 7.1 for insertions, and 5.8 for deletions. The horizontal dashed line indicates a ratio of 1.

(E) Fraction of CSA-SVs and other CDH-overlapping SVs located within the indicated distance of a TAD boundary or chromatin-loop anchor. Lines show medians and shaded regions indicate IQRs. At a distance threshold of 25 kbp, a median of 28.4% of CSA-SVs overlapped or occurred near one of these chromatin architectural features.

(F) Genomic annotation of CSA-SVs according to overlap with protein-coding exons. Of 2,495 CSA-SVs, 2,478 (99.3%) did not overlap a protein-coding exon, whereas 17 (0.7%) overlapped a protein-coding exon.

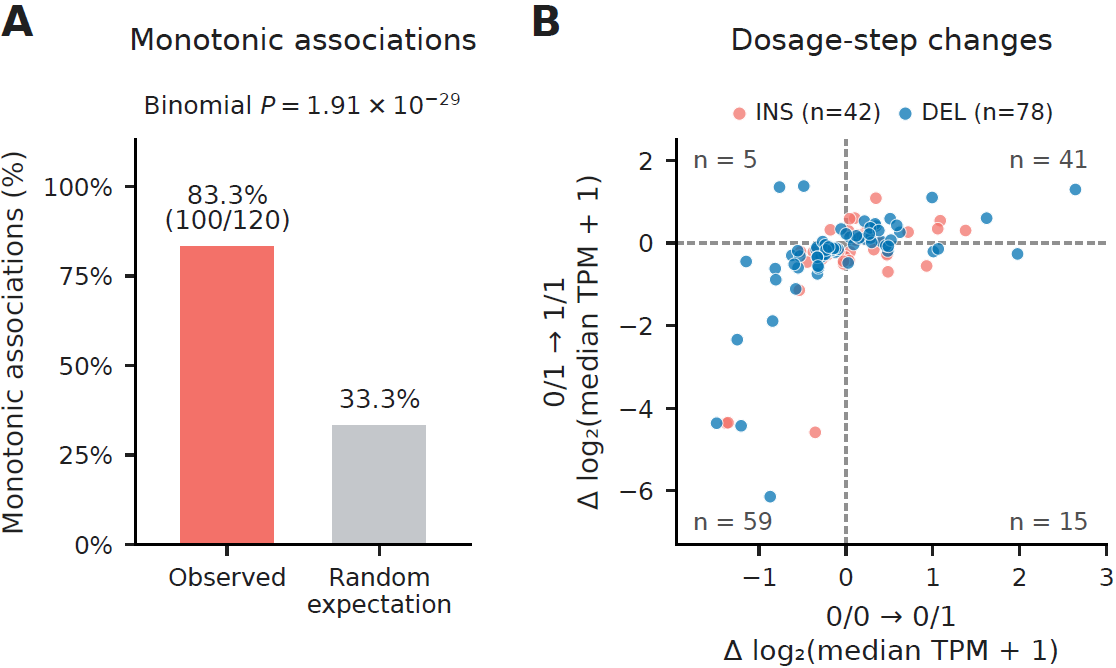

**Figure S8. Genotype-associated expression patterns for CSA-SV**–**gene associations, related to Figure 4**

(A) Observed proportion of monotonic expression patterns among 120 significant CSA-SV–gene associations with all three genotype classes represented and distinct genotype-class median expression levels. Of these associations, 100 (83.3%) showed monotonically increasing or decreasing expression across 0/0, 0/1, and 1/1 genotypes, compared with the 33.3% expectation under random ordering of three distinct medians (*P* = 1.91 × 10⁻²⁹, two-sided binomial test).

(B) Expression changes between consecutive genotype classes for the same 120 associations. The horizontal and vertical axes show changes in log₂(median TPM + 1) from 0/0 to 0/1 and from 0/1 to 1/1, respectively. Each point represents one CSA-SV–gene association, colored by SV type: insertions (coral; n = 42 associations) and deletions (blue; n = 78 associations). Dashed lines indicate zero change, and numbers indicate the association count in each quadrant. The upper-right and lower-left quadrants represent monotonically increasing and decreasing expression patterns, respectively.

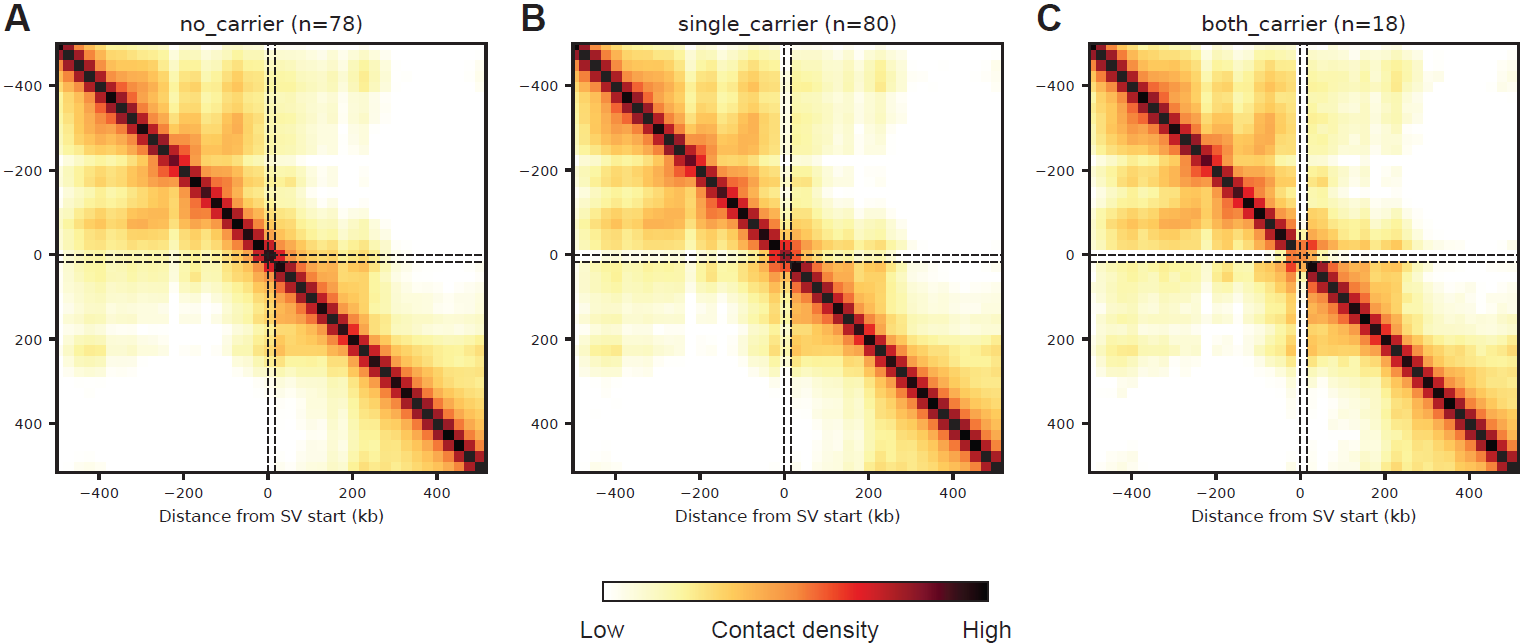

**Figure S9. Genotype-stratified chromatin contact maps at the *SESTD1*-associated CSA-SV locus, related to Figure 4.**

(A–C) Mean normalized chromatin contact maps surrounding the 16.2 kbp deletion chr2-179683834-DEL-16208 in non-carriers (0/0; A, n = 78), heterozygous carriers (0/1; B, n = 80), and homozygous deletion carriers (1/1; C, n = 18). Contact matrices were extracted at 25 kbp resolution across the SV interval and its 500 kbp flanking regions. Each individual’s matrix was normalized by its total contact signal within the extracted region before averaging across individuals within each genotype class. Both axes indicate distance from the SV start. Dashed lines mark the SV start and end on both axes. Colors indicate mean normalized contact density on a shared logarithmic scale.

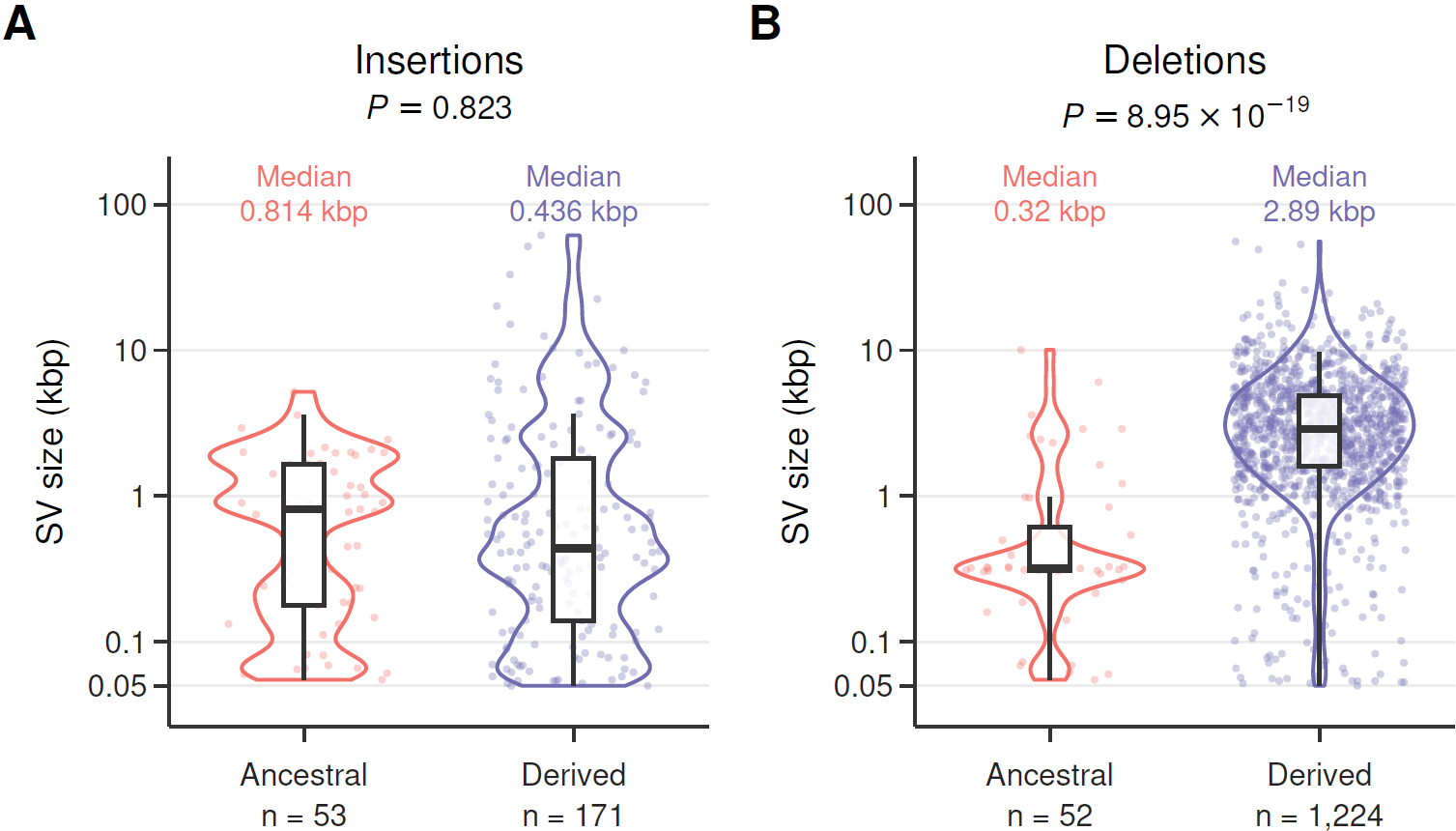

**Figure S10. Size distributions of ancestral and human-derived CSA-SVs by variant type, related to Figure 5.**

(A) Size distributions of ancestral (n = 53) and human-derived (n = 171) CSA-SV insertions (medians, 0.814 and 0.436 kbp, respectively; *P* = 0.823).

(B) Size distributions of ancestral (n = 52) and human-derived (n = 1,224) CSA-SV deletions (medians, 0.320 and 2.89 kbp, respectively; *P* = 8.95 × 10⁻¹⁹).

Ancestral and human-derived variants are shown in coral and purple, respectively. Violin outlines show the distributions, and individual points represent CSA-SVs. Boxes indicate IQRs, horizontal lines within boxes indicate medians, and whiskers extend to the most extreme observations within 1.5 times the IQR below the first quartile or above the third quartile. *P* values were calculated using two-sided Wilcoxon rank-sum tests with continuity correction.

### SUPPLEMENTAL TABLES

Supplemental Tables S1-S11 provided in separate Excel file.

**Table S1. Sample information and chromatin contact sequencing yields, related to Figure 1**

Sample metadata and sequencing yields for the 177 HPRC Release 2 individuals included in this study. Columns report sample identifiers, population abbreviations and descriptions, total sequencing yield, chromatin contact assay, and the reported number of replicates. Each row represents one individual.

**Table S2. Read-pair statistics for T2T-CHM13 mappings, related to Figure 1**

Read-pair processing statistics for the 177 individuals after mapping to T2T-CHM13. Counts are reported for total, mapped, PCR-duplicate, and nonduplicate read pairs. Each row represents one individual. No-Dup, nonduplicate.

**Table S3. Read-pair statistics for DSA mappings, related to Figure 1**

Total, mapped, and nonduplicate valid read-pair counts after independently mapping each individual’s data to the corresponding Hap1 and Hap2. The same input read pairs were mapped independently to both assemblies; the two total-read columns therefore represent the same sequencing input.

**Table S4. Reference-dependent changes in inferred contact distance, related to Figure 1**

Per-haplotype counts of shared contact pairs used to compare inferred genomic distances between DSAs and T2T-CHM13. Long5, Long10, Long25, Long50, and Long100 report counts at shift thresholds of 5, 10, 25, 50, and 100 kbp, respectively. Cis gives the number of retained shared *cis* contact pairs used as the denominator for shift fractions; Shared gives the total shared contact-pair count. Each row represents one individual haplotype comparison.

**Table S5. Haplotype-specific contact difference hotspot windows, related to Figure 2**

Catalog of CDH windows across the haplotypes. Each row specifies a window’s chromosome, start and end coordinates, haplotype, and sample identifier. Coordinates are given in the corresponding DSA, with chromosome scaffold names retaining the RagTag suffix.

**Table S6. Haplotype-specific counts of heterozygous SVs, related to Figure 2**

Counts of heterozygous SVs used for CDH–SV overlap analyses. Counts are reported separately for insertions and deletions for each individual and haplotype. SV_count gives the number of variants in the indicated category.

**Table S7. Recurrent CDH regions in T2T-CHM13 coordinates, related to Figure 3**

Catalog of recurrent CDH regions identified after projecting haplotype-specific CDHs to T2T-CHM13. Columns provide chromosome, start and end coordinates, region identifier, region length, and the minimum and maximum positional haplotype support within each region.

**Table S8. SVs associated with local CDHs, related to Figure 3**

Catalog of CSA-SVs. Each row reports an SV identifier, type and length, its position in T2T-CHM13, the flanking distance used to assess CDH overlap, haplotype contingency counts, an OR, and raw and adjusted *P* values.

**Table S9. Long-read transcriptome sequencing yields, related to Figure 4**

Numbers of FLNC RNA-seq reads for the 163 individuals with matched chromatin contact data used in the expression analyses. Each row reports a sample identifier and its FLNC read count after long-read RNA-seq preprocessing.

**Table S10. Significant associations between CSA-SV genotypes and gene expression, related to Figure 4**

Summary of significant *cis* associations between CSA-SVs and expression of genes. Each row represents one SV–gene pair selected by a Kruskal–Wallis test with a Benjamini–Hochberg-adjusted *P* value below 0.05 (q_kw). n_total and n_groups give the number of individuals with valid observations and the number of represented genotype classes, respectively. AA, Aa, and aa denote genotypes 0/0, 0/1, and 1/1, corresponding to zero, one, and two copies of the alternate allele. n_AA, n_Aa, and n_aa give the genotype-specific sample counts; med_AA, med_Aa, and med_aa give median expression in TPM. Blank median entries indicate an unrepresented genotype class. p_kw is the raw Kruskal–Wallis *P* value. rho is the Spearman correlation between alternate-allele dosage and TPM, with p_trend and q_trend giving its raw and Benjamini–Hochberg-adjusted two-sided *P* values. svtype and svlen give SV type and length.

**Table S11. Inferred evolutionary origins of CSA-SV alleles, related to Figure 5**

Evolutionary-origin annotations for all CSA-SVs. Columns report chromosome, variant identifier, SV type, SV length in base pairs, presence of the variant record in the ape genotype VCF (VCF_PRESENT), and inferred origin. VCF_PRESENT is a record-availability flag, with TRUE and FALSE denoting presence and absence of a record, respectively. VCF, variant call format.
